# The Manifold Actions of Signaling Peptides on Subcellular Dynamics of a Receptor Specify Stomatal Cell Fate

**DOI:** 10.1101/836437

**Authors:** Xingyun Qi, Michal Maes, Scott M. Zeng, Keiko U. Torii

**Affiliations:** Howard Hughes Medical Institute and Department of Biology, University of Washington, Seattle, WA 98195, USA; Department of Physics, University of Washington, Seattle, WA 98195, USA; Howard Hughes Medical Institute and Department of Molecular Biosciences, University of Texas at Austin, Austin, TX 78712, USA

## Abstract

Receptor endocytosis is important for signal activation and transduction. However, how a receptor interprets conflicting signals to adjust cellular output is not clearly understood. During plant development, the family of EPIDERMAL PATTERNING FACTOR (EPF) peptides fine-tunes stomatal patterning through ERECTA-family receptor kinases. Using genetic, cell biological, and pharmacological approaches, we report here that ERECTA-LIKE1 (ERL1), the major receptor restricting stomatal differentiation, undergoes dynamic subcellular behaviors in response to different signal inputs. ERL1 is constitutively recycled, whereas its activation by EPF1 peptide induces rapid internalization to multivesicular bodies (MVB). In contrast, dominant-negative ERL1 resides predominantly in plasma membrane. The co-receptor, TOO MANY MOUTHS (TMM), is essential for EPF1-induced ERL1 internalization but dispensable for EPFL6-induced ERL1 internalization. The peptide antagonist of EPF1, Stomagen/EPFL9, triggers retention of ERL1 in the endoplasmic reticulum. Our study elucidates that multiple related yet unique peptides specify cell fate by deploying the differential subcellular dynamics of a single receptor.

## Introduction

Receptor-mediated endocytosis is an integral part of cellular signaling, as it mediates signal attenuation and provides spatial and temporal dimensions to signaling events. In mammalian systems, endocytosis of receptor tyrosine kinases can attenuate the signal outputs, by removing the active receptor pools from the plasma membrane, or it can specify signals at defined sites of action, such as signaling through endosomes (Sigismund et al., 2012). As a sessile organism, plants make use of a large number of receptor-like kinases (RLKs) for cell-cell, shoot-to-root, and inter-kingdom communications (Shiu and Bleecker, 2001). The RLKs with extracellular leucine-rich repeat domain, known as LRR-RLKs, comprise the largest RLK subfamily (Shiu and Bleecker, 2001), and they specify critical aspects of development, environmental response, and immunity by perceiving extrinsic signals (Torii, 2004; Macho and Zipfel, 2014). Increasing evidence shows that the subcellular localization and trafficking routes of LRR-RLKs regulate their function and activity (Ben Khaled et al., 2015). In Arabidopsis, bacterial flagellin peptide flg22 induces the heterodimer formation consisting of the LRR-RLKs FLAGELLIN SENSING2 (FLS2) and BRI1-ASSOCIATED RECEPTOR KINASE (BAK1)/SOMATIC EMBRYOGENESIS RECEPTOR LIKE KINASE 3 (Chinchilla et al., 2007). This triggers the endocytosis and degradation of the complex to generate transient cellular immune signaling but also to prevent continuous signaling to the same stimulus (Robatzek et al., 2006; Beck et al., 2012). The brassinosteroid (BR) receptor BRASSINOSTEROID INSENSITIVE1 (BRI1) forms a complex with BAK1 (Li et al., 2002; Nam and Li, 2002; Bücherl et al., 2013). BRI1 can undergo constitutive endocytosis independent of BRs, but BRs can elevate BRI1 and BAK1 interaction and reduce the number of available BRI1-BAK1 complexes on the plasma membrane (Geldner et al., 2007; Bücherl et al., 2013; Hutten et al., 2017). CLAVATA1 (CLV1), an LRR-RLK that controls stem cell homeostasis within the shoot meristem (Clark et al., 1997), is downregulated by ligand-dependent internalization upon perception of its ligand CLV3 to buffer the signal of CLV3 (Nimchuk et al., 2011). It remains a key question as to where within the cell these LRR-RLKs transduce signals and how different activation states of LRR-RLKs influence their subcellular localization.

Developmental patterning of stomata, adjustable pores on the plant epidermis for gas-exchange and transpiration, relies on intricate cell-cell communication mediated by signaling peptides and their receptors (Lau and Bergmann, 2012; Pillitteri and Torii, 2012). In Arabidopsis, secreted peptides from the EPF family, and their shared receptor LRR-RLKs ERECTA, ERL1 and ERL2, mediate this process (Rychel et al., 2010). Amongst the plant LRR-RLKs, the ERECTA family offers a unique advantage to study how multiple signals are perceived to achieve cell fate and patterning. EPF2 and EPF1 negatively regulate stomatal development primarily through ERECTA and ERL1, respectively (Hara et al., 2007; Hara et al., 2009; Hunt and Gray, 2009). In contrast, EPF-LIKE9 (EPFL9), also known as Stomagen, promotes stomatal development by competing with EPF2 and, to some extent, with EPF1 for receptor binding (Sugano et al., 2010; Lee et al., 2015; Lin et al., 2017; Qi et al., 2017). Moreover, EPFL4/5/6, a subfamily only expressed in hypocotyls and stems, also act as ligands for the ERECTA family to inhibit stomatal formation when an LRR receptor protein, TMM, is missing (Abrash and Bergmann, 2010; Abrash et al., 2011). Although the final phenotypic outcomes of these different EPF signaling events are well characterized, the very early step of signal transmission by the receptors remain elusive. While internalization of ERL2 was documented briefly (Ho et al., 2016), it is unknown whether it has any implications in signal transduction or in which subcellular organelle ERL2 was localized.

Among the ERECTA family, ERL1 regulates guard cell differentiation in an autocrine manner in addition to enforcing stomatal spacing of neighboring cells in a paracrine manner (Lee et al., 2012; Qi et al., 2017). This dual function of ERL1 can be attributed to its cell-type specific expression patterns as well as its ability to perceive different EPF/EPFL peptide ligands (Shpak et al., 2005; Lin et al., 2017). It remains unknown, however, how ERL1 receptor dynamics translate into the eventual stomatal cell fate. Here, we combined genetic, pharmacological, and live imaging approaches to explore the initial events that occurred at ERL1 upon perception of different EPF peptides. Our study shows that EPF1 and EPFL6, the ligands activating the inhibitory stomatal signaling, trigger ERL1 endocytosis into MVBs. TMM, which can form a receptor complex with ERL1, is required for the EPF1-induced ERL1 internalization and suppression of stomatal fate, but is superfluous for EFPL6-induced ERL1 internalization. Surprisingly, Stomagen interferes with the inhibitory regulation of stomatal differentiation by retaining ERL1 to the endoplasmic reticulum, similar to when endocytosis was pharmacologically blocked by Tyrphostin A23 (Tyr A23) (Santuari et al., 2011) and Endosidin 9-17 (ES9-17) (Dejonghe et al., 2019). Our study reveals a mechanism by which plant cells interpret multiple signals through the subcellular localization and trafficking route of a single receptor.

## Results

### ERL1 is internalized through multivesicular bodies to vacuolar pathway in stomatal meristemoids

To understand how stomatal cell fate decisions are made at the level of receptor subcellular dynamics, we first examined the localization of ERL1 (Figure 1). As reported previously (Qi et al., 2017), a functional ERL1-YFP fusion protein driven by its endogenous promoter (ERL1pro::ERL1-YFP) in *erl1* seedlings marks the plasma membrane of stomatal-lineage cells, most notably differentiating meristemoids. In addition, we detected some highly mobile punctae highlighted by ERL1-YFP within the cells (Figure 1A, Video 1). To define the subcellular localization of ERL1-YFP, its co-localization analysis was performed with marker proteins Syp43-RFP for trans-Golgi network (TGN), RFP-Ara7 for MVB, and Syp22-RFP for vacuoles (occasionally MVB) (Figure 1A). ERL1-YFP extensively co-localizes and moves together with RFP-Ara7 (Figure 1A, B, Video 1), whereas only 25 % and 18 % of ERL1-YFP-positive punctae are also labelled by Syp43-RFP and Syp22-RFP, respectively. Thus, ERL1-YFP predominantly resides on the MVB. This is further confirmed by a pharmacological approach using Wortmannin (Wm), a fungal drug that can cause fusion of MVBs by inhibiting phosphatidylinositol-3 (PI3) and phosphatidylinositol-4 (PI4) kinases (Foissner et al., 2016). The Wm application on Arabidopsis seedlings resulted in the formation of typical ring-like Wm bodies marked by both ERL1-YFP and RFP-Ara7 (Figure 1C). Taken together, these results indicate that, within the stomatal precursor cells, ERL1 undergoes endocytic trafficking from plasma membrane to MVB.

**Figure 1.**
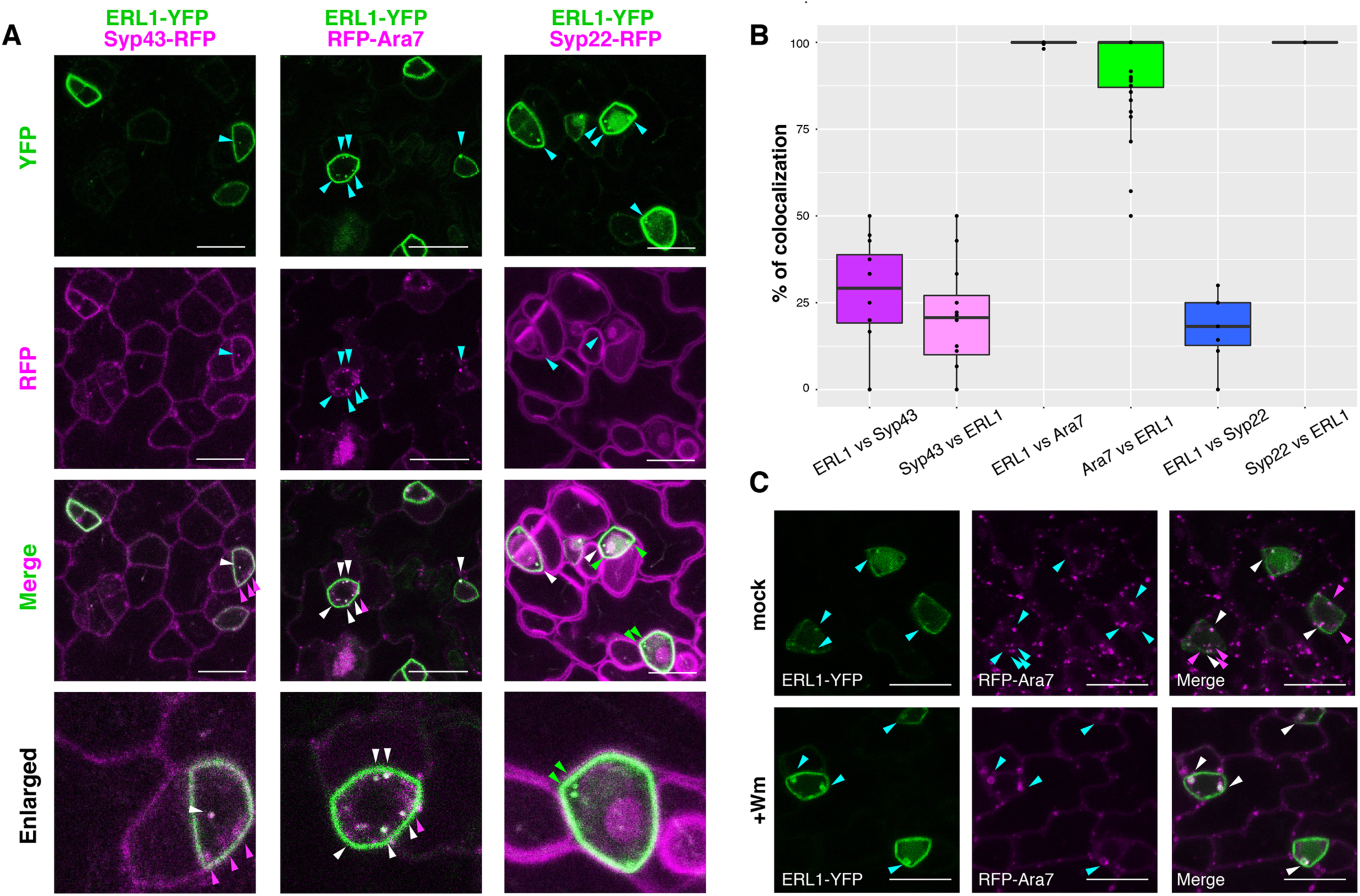
ERL1-YFP has dual localization on plasma membrane and late endosomes. (**A**) Representative confocal microscope images of ERL1-YFP (top row) co-localization analysis with the TGN marker Syp43-RFP (left column), the MVB marker RFP-Ara7 (middle column), and the MVB and vacuole marker Syp22-RFP (right column) in the abaxial epidermis of developing true leaves of the 7-day-old seedlings. Merged images are shown in the third row, with enlarged images of representative meristemoids in the bottom row. Arrowheads point to endosomes bearing ERL1-YFP, RFP-Syp43, RFP-Ara7, and/or RFP-Syp22: cyan, single channels (top two rows); green, YFP; magenta, RFP; white, co-localization (bottom two rows). Scale bars = 10 µm. (**B**) Quantitative analysis of the co-localized endosomes between ERL1-YFP and the subcellular marker proteins. Percentage of the endosomes of the former protein that co-localize with the latter protein is shown as dots. Lines in the boxplot show the median value of each group, and the boxes represent from the first to third quartiles. n = 40 for ERL1 vs Ara7 or Ara7 vs ERL1; n = 12 for ERL1 vs Syp43 or Syp43 vs ERL1; n=7 for ERL1 vs Syp22 or Syp22 vs ERL1. (**C**) ERL1-YFP and RFP-Ara7 treated with Wm. Shown are ERL1-YFP (left column) and RFP-Ara7 (middle column) in the abaxial epidermis of developing true leaves of the 7-day-old seedlings treated with mock (top row) or 30 µM Wm (bottom row). Arrowheads point to ERL1-YFP and/or RFP-Ara7 endosomes: cyan, single channels; magenta, YFP; white, co-localization. Scale bars = 10 µm.

### TMM is required for the process of ERL1 endocytosis in true leaves

Endocytosis is an essential process to regulate cell signaling by controlling the turnover of plasma membrane proteome. We wondered if ERL1 endocytosis is related to its biological signaling. A previous work has shown that ERL1 forms a heterodimer with TMM, a receptor protein, to create a pocket for the proper binding of its major ligand EPF1 (Lee et al., 2012; Lin et al., 2017). The absence of TMM results in clustered stomata (Figure 2A), indicating that TMM is required for EPF1-ERL1 signaling to enforce proper stomatal spacing (Hara et al., 2007; Lee et al., 2012). As a first step to test whether active ERL1 signaling is a prerequisite for its endocytosis, we monitored ERL1 dynamics in *tmm* background (Figure 2A). Interestingly, the number of cells with ERL1-YFP-positive endosomes is greatly reduced in *tmm* mutant (63% in WT (n=323 cells) *vs*. 30% in *tmm* (n=466 cells)).

**Figure 2.**
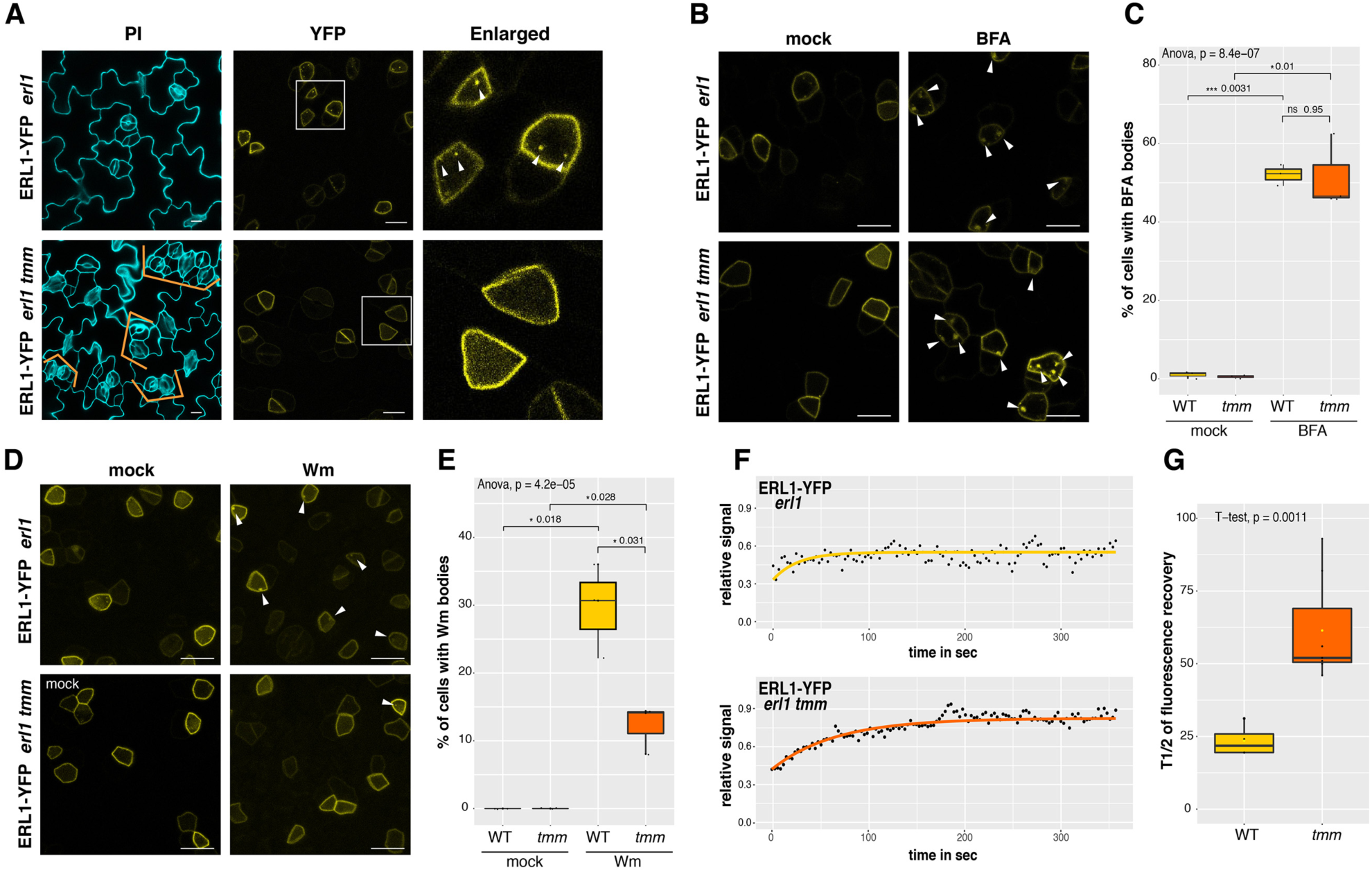
ERL1 internalization requires its co-receptor TMM. (**A**) Representative confocal microscopy images of ERL1-YFP in *erl1* (top row) and in *erl1 tmm* (bottom row) in the abaxial epidermis of developing true leaves of the 7-day-old seedlings. Right column; enlarged images. Their stomatal phenotypes are shown on the left column. Orange brackets: clustered stomata. Arrowheads indicate endosomes. Scale bars =10 µm. (**B**) Representative images of ERL1-YFP in *erl1* (top row) or in *erl1 tmm* (bottom row) of the abaxial epidermis of developing true leaves from the 7-day-old seedlings treated with mock (left column) or 30 µM BFA (right column). Arrowheads indicate BFA bodies. Scale bars =10 µm. (**C**) Quantitative analysis of the number of cells with BFA bodies when ERL1-YFP in *erl1* (yellow) or *erl1 tmm* (orange) are treated with mock or 30 µM BFA. Lines in the boxplot show the median value. ANOVA was performed for comparing all samples, and Student’s T-test was performed for pairwise comparisons. ns, not significant. * p<0.05. *** p<0.0005 n=3 independent experiments. For each experiment, the total numbers of cells counted are 86, 130, 66 (WT mock); 138, 94, 131 (*tmm* mock); 230, 336, 109 (WT BFA); 300, 393, 211 (*tmm* BFA). (**D**) Representative images of ERL1-YFP in *erl1* (top row) or in *erl1 tmm* (bottom row) treated with mock (left column) or 25 µM Wm (right column). Arrowheads indicate Wm bodies. Scale bars =10 µm. (**E**) Quantitative analysis of the number of cells with Wm bodies when ERL1-YFP in *erl1* (yellow) or *erl1 tmm* (orange) are treated with mock or 25 µM Wm. Lines in the boxplot show the median value. ANOVA was performed for comparing all samples, and Student’s T-test was performed for pairwise comparisons between *erl1* and *erl1 tmm*. * p<0.05. *** p<0.0005 n=3 independent experiments. For each experiment, the total numbers of cells counted are 155, 73, 62 (WT mock); 181, 126, 104 (*tmm* mock); 184, 136, 176 (WT Wm); 1043, 273, 83 (*tmm* BFA). (**F**) FRAP analyses of plasma membrane ERL1-YFP in wild type (*erl1-2*) or in *tmm* (*erl1-2 tmm*). Shown are representative fluorescence recovery curves plotted as a function of time and fitted to Single Exponential Fitting. ERL1-YFP in *erl1* (top; yellow); ERL1-YFP in *erl1 tmm* (bottom; orange). (**G**) Quantitative analysis of the half time of fluorescence recovery of plasma membrane ERL1-YFP in *erl1* (yellow) and *erl1 tmm* (orange). Lines in the boxplot show the median value. T-test was performed for pairwise comparisons between *erl1* and *erl1 tmm*. n=3 for WT and n=9 for *tmm*.

Activated plant receptor kinases can either be recycled back to the plasma membrane or are destined for endocytic degradation via MVB for signaling termination. To address whether TMM is required for a specific pathway, we first treated Arabidopsis *ERL1pro::ERL1-YFP* seedlings in wild type (*erl1*) and *tmm* (*erl1 tmm*) background with a membrane-trafficking drug brefeldin A (BFA), a chemical inhibitor of GNOM, an ADP-ribosylation factor - guanine nucleotide exchange factor that mediates endosomal recycling (Geldner et al., 2003). When treated with BFA, ERL1-YFP-positive BFA bodies were detected in both wild type and *tmm* mutant background with no significant difference (Figures 2A, C). Furthermore, BFA treatment in the presence of protein synthesis inhibitor cycloheximide (CHX) conferred ERL1-YFP-positive BFA body formation in both wild type and *tmm* mutant with no discernable difference (Figure S1). Thus, the results indicate that ERL1 proteins deriving from an endosomal recycling pathway, but not from a secretory pathway, contribute to BFA body formation and that TMM does not influence recycling of ERL1.

Next, the seedlings were treated with Wm. In sharp contrast to the BFA treatment, the Wm treatment conferred significant reduction of ERL1-YFP-marked Wm-bodies in *tmm* compared to that in wild type (Figure 2D and E, 30% in wild type *vs*. 12% in *tmm*, p = 0.031, Student’s t-test). Combined, the results suggest that TMM is essential for the internalization of ERL1 to MVB, rather than the recycling of ERL1 to the plasma membrane. To rule out the possibility that the reduced ERL1 endocytosis in *tmm* is due to defects in the general endocytic degradation machinery, we examined the effects of *tmm* on general endocytosis using FM4-64, a styryl dye used to trace the endocytic pathways in Arabidopsis (Meckel et al., 2004)(Figure S1A). In wild type, 92.8% (n=20 cells) of ERL1-YFP-labelled endosomes can be stained by FM4-64. In *tmm*, however, FM4-64 still internalizes to multiple endosomes whereas ERL1-YFP fails to internalize in 70% of the cells examined (n=30 cells) (Figure S1A). We next examined the effects of *tmm* on the formation of MVBs. In the cells co-expressing RFP-Ara7 and ERL1-YFP, no significant difference was observed in the numbers of RFP-Ara7-marked endosomes and Wm bodies between wild type and *tmm* (3.68 endosomes/cell in wild type *vs*. 4.12 endosomes/cell in *tmm* and 2.45 Wm bodies/cell in wild type *vs*. 3.00 Wm bodies/cell in *tmm*). In contrast, the *tmm* mutation conferred substantial reduction in ERL1-YFP-marked endosomes and Wm bodies (2.27 endosomes/cell in wild type *vs*. 0.78 endosomes/cell in *tmm* and 1.84 Wm bodies/cell in wild type *vs*. 0.73 Wm bodies/cell in *tmm*), all of which colocalized with RFP-Ara7 (Figure S2B, C). Thus, TMM is specifically required for ERL1’s endocytic sorting pathway to MVB, a hallmark for eventual receptor degradation in a vacuole (Geldner and Robatzek, 2008).

To further explore the role of TMM for the ERL1 receptor dynamics on plasma membrane, we performed fluorescence recovery after photobleaching (FRAP) assays on ERL1-YFP on the plasma membrane, and the half time of fluorescence recovery was calculated from modeling to exponential curves (Figure 2F and G). In wild type, the calculated mean half time of ERL1-YFP fluorescence recovery (t1/2) was 23.55 ± 5.55 sec, whereas in *tmm* it was 70.89 ± 24.63 sec (Figure 2G). The longer recovery time of ERL1-YFP in *tmm* could be explained by the slower removal of the photobleached receptor molecules from the plasma membrane due to decreased internalization. Combined, these results support a notion that, in the absence of TMM, un-activated ERL1 receptors are not readily targeted for endocytic pathway and, consequently, remain stable on the plasma membrane.

### Dominant-negative ERL1 receptor is predominantly at the plasma membrane

It has been shown that removal of the cytoplasmic kinase domain from ERECTA-family RLKs confers strong dominant-negative effects both in aboveground organ growth and in stomatal patterning (Shpak et al., 2003; Lee et al., 2012). The dominant-negative ERL1ΔK can directly binds its ligand EPF1 through the extracellular LRR domain. However, it is unable to signal and, consequently, confers paired and clustered stomata, thereby phenocopying *epf1* mutant (Figure 3A and B) (Lee et al., 2012). We examined the subcellular dynamics of the dominant-negative ERL receptor, ERL1ΔK fused with CFP driven by its endogenous promoter (*ERL1pro::ERL1*Δ*K-CFP*) (Figure 3C). Strong CFP signal is detected on the plasma membrane of stomatal precursor cells, but only very few mobile punctae can be seen within cells (Figure 3C). Similar to ERL1 behavior in *tmm* mutant, the dominant-negative ERL1 is sensitive to BFA treatment (Figure 3D, F and G), with 86% cells possessing ERL1ΔK-CFP-marked BFA bodies. This BFA sensitivity of ERL1ΔK was also observed in the presence of CHX (Figure S1), indicating that they represent recycling populations. In contrast, ERL1ΔK-CFP exhibits insensitivity to Wm treatment, with only 18% cells showing Wm bodies highlighted by ERL1ΔK-CFP (Figure 3D, F and G). Notably, the reduced endocytosis of ERL1ΔK-CFP is not due to defects in the general endocytosis process, as FM4-64 can still internalize in the ERL1ΔK-CFP positive cells on the transgenic seedling epidermis, like it does in cells with the full-length ERL1 (Figure 3E). These results suggest that activation of ERL1 signaling is required for the receptor internalization.

**Figure 3.**
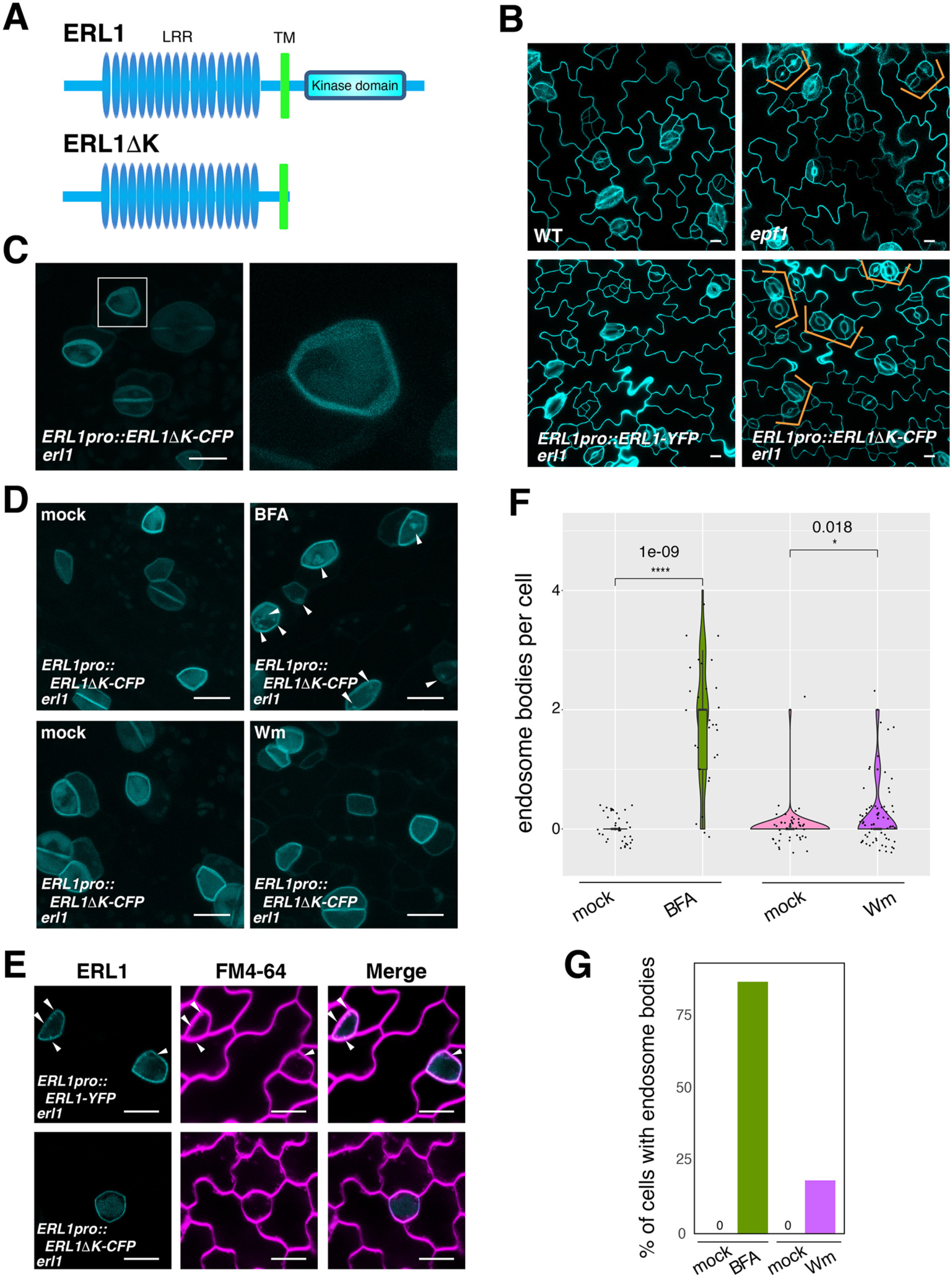
ERL1 internalization requires its functionality. (**A**) Diagram of the full-length ERL1 protein (top) and the dominant-negative ERL1 protein lacking the cytoplasmic domain (bottom). (**B**) Representative confocal microscopy images of cotyledon abaxial epidermis from the 4-day-old seedlings of wild type (top left), *epf1* (top right), ERL1-YFP *erl1* (bottom left) and ERL1ΔK-CFP *erl1* (bottom right), stained by PI. Brackets indicate the paired stomata in *epf1* and ERL1ΔK-CFP in *erl1*. Scale bars =10 µm. (**C**) Representative confocal microscopy images of ERL1ΔK-CFP in *erl1* of the abaxial developing true leaf epidermis from the 7-day-old seedlings. Right, the enlarged image from the highlighted area (left, white rectangle). Scale bars =10 µm. (**D**) Representative confocal microscopy images of ERL1ΔK-CFP treated with mock (top left for BFA treatment), 30 µM BFA (top right), mock (bottom left for Wm treatment) and 25 µM Wm (bottom right). Arrowheads indicate BFA bodies. Scale bars =10 µm. (**E**) Representative images of ERL1-YFP in *erl1* (upper row) and ERL1ΔK-CFP in *erl1* (bottom row) stained with an endocytosis monitoring membrane dye, FM4-64, in the abaxial epidermis of developing true leaves of the 7-day-old seedlings. Arrowheads indicate internalized endosomes. (**F**) Quantitative analysis of the number of ERL1-YFP-positive BFA- or Wm bodies per cell shown as a violin plot. Individual data points are dot-plotted with jitter. Median values are shown as lines in the boxplot. ANOVA was performed for comparing all samples, and T-test was performed for the pairwise comparison of mock and drug-treated samples. p values were indicated between every two compared samples. n = 36 for mock (BFA); n = 29 for BFA; n = 44 for mock (Wm); n = 66 for Wm. (**G**) Quantitative analysis of the percentage of cells with BFA bodies (green) or Wm bodies (purple) when ERL1ΔK-CFP are treated with mock, 30 µM BFA or mock, 25 µM Wm.

### EPF1 triggers TMM-dependent ERL1 internalization

Of the 11 EPF family members, EPF1 is the major ligand for ERL1 (Lee et al., 2012). EPF1 signaling plays a negative role in stomatal development, and the induction of EPF1 peptide (iEPF1) confers arrested stomatal precursors (Figure 4A) (Hara et al., 2007; Lee et al., 2012; Qi et al., 2017). We therefore tested whether the ERL1 internalization is ligand dependent. For this purpose, we first examined ERL1-YFP dynamics in *epf1* mutants. As shown in Figure S3A, both plasma membrane and highly mobile endosomes are highlighted by ERL1-YFP in *epf1*. When treated with BFA or Wm, the percentages of cells with ERL1-YFP-marked BFA or Wm bodies are similar between wild type and *epf1* (Figure S3B-E), indicating the general trafficking of ERL1-YFP is not severely affected in the absence of EPF1.

**Figure 4.**
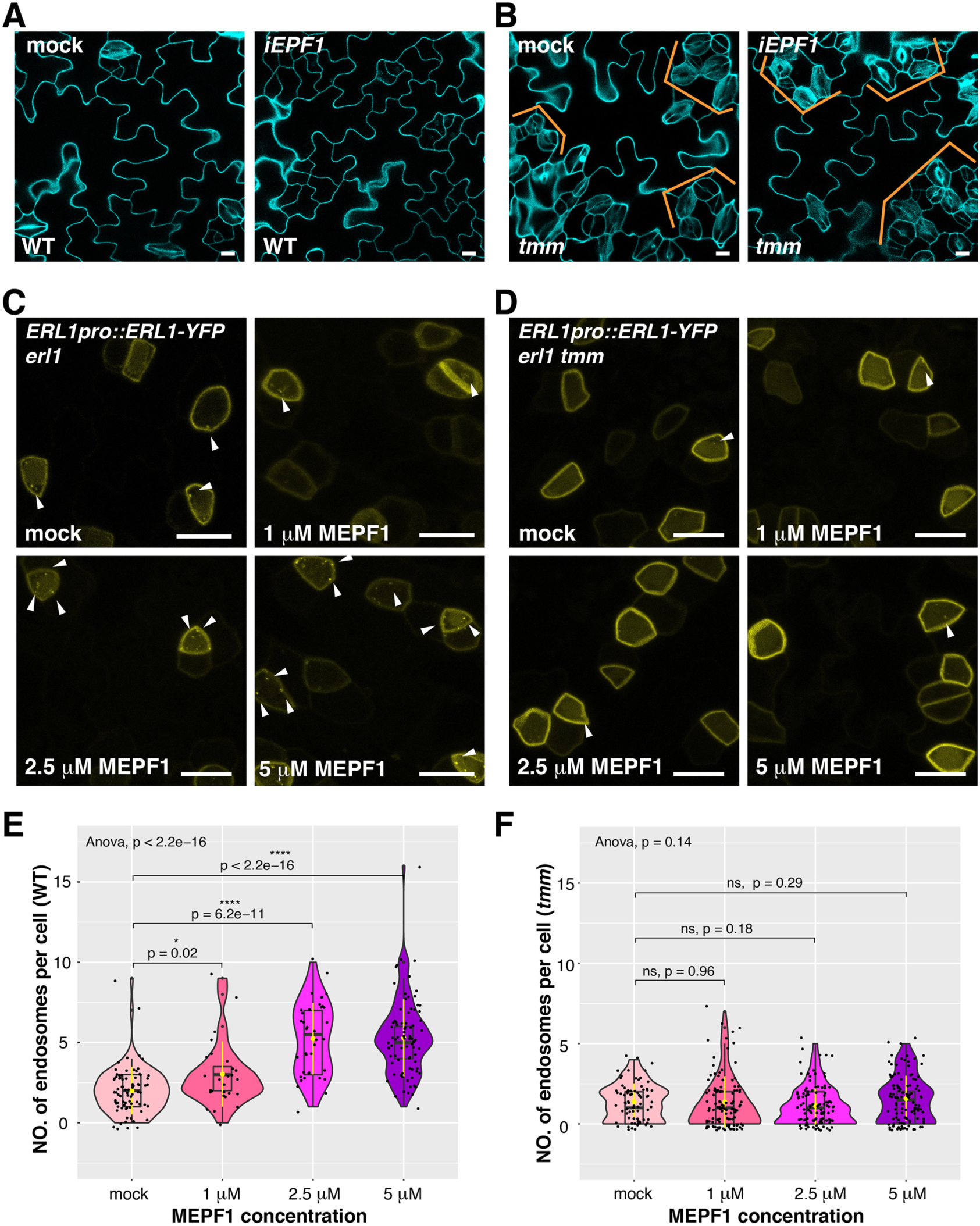
MEPF1 triggers ERL1-YFP internalization in *erl1* but not in *erl1 tmm*. (**A**) Representative confocal microscopy images of cotyledon abaxial epidermis from the 4-day-old *iEPF1* seedlings treated with mock (left) or 10 µM Estradiol (right). Scale bars =10 µm. (**B**) Representative confocal microscopy images of cotyledon abaxial epidermis from the 4-day-old *iEPF1* in *tmm* seedlings treated with mock/DMSO (left) or 10 µM Estradiol (right). Brackets indicate clustered stomata in both mock- and estradiol-induced samples. Scale bars =10 µm. (**C**) Representative confocal microscopy images of ERL1-YFP in *erl1* treated with mock (top left), 1 µM MEPF1 (top right), 2.5 µM MEPF1 (bottom left) and 5 µM MEPF1 (bottom right) are shown. Arrowheads indicate endosomes. Scale bars = 10 µm. (**D**) Representative confocal microscopy images of ERL1-YFP in *erl1 tmm* treated with mock (top left), 1 µM MEPF1 (top right), 2.5 µM MEPF1 (bottom left) and 5 µM MEPF1 (bottom right) are shown. Arrowheads indicate endosomes. Scale bars = 10 µm. (**E**) Quantitative analysis of the number of ERL1-YFP-positive endosomes per cell at different concentrations of MEPF1 application in *erl1* shown as a violin plot. Dots, individual data points. Median values are shown as lines in the boxplot, and mean values are shown as yellow dots in the plot. ANOVA was performed for comparing all samples, and T-test was performed for pairwise comparisons of samples treated with the mock and different concentration of MEPF1. n= 79, 27, 38, 82 for treatment with mock, 1 µM, 2.5 µM, 5µM MEPF1. (**F**) Quantitative analysis of the number of ERL1-YFP-positive endosomes per cell at different concentrations of MEPF1 application in *erl1 tmm* shown as a violin plot. Dots, individual data points. Median values are shown as lines in the boxplot, and mean values are shown as yellow dots in the plot. ANOVA was performed for comparing all samples, and T-test was performed for pairwise comparisons of samples treated with the mock and different concentration of MEPF1. n= 76, 113, 109, 114 for treatment with mock, 1 µM, 2.5 µM, and 5µM MEPF1, respectively.

Considering the high similarity among the 11 EPF members, it is possible that the functional redundancy of other EPFs alleviates the defect of ERL1 internalization in *epf1*. To overcome the genetic redundancy, we took advantage of the biologically-active mature EPF1 (MEPF1) peptide (Figure 4) (Lee et al., 2012; Qi et al., 2017). Different concentrations of MEPF1 were applied to the true leaf epidermis of 7-day-old seedlings expressing *ERL1pro::ERL1-YFP*. The number of ERL1-YFP-positive endosomes per cell increases as the peptide concentration increases (Pearson correlation, r=0.56, p= 2.2 e-16; Figure S3C, E), indicating that MEPF1 peptide triggers the internalization of ERL1 in a dosage-dependent manner. In the *tmm* background, however, the number of ERL1-YFP-positive endosomes per cell remains low regardless of the MEPF1 dosage applied (Figure 4D, F). Thus, in the absence of TMM, ERL1-YFP endocytosis is insensitive to MEPF1 application, consistent with the genetic evidence that the *tmm* mutation is epistatic to induced *EPF1* overexpression (*iEPF1*) (Figure 4B) (Hara et al., 2007; Lee et al., 2012). Taken together, we conclude that EPF1 peptide ligand perception triggers the internalization of ERL1 receptor in a TMM-dependent manner.

### EPFL6 triggers ERL1 internalization in the absence of TMM

A previous structural analysis has shown that binding of EPF1 to the ERL1-TMM receptor complex does not lead to conformational change (Lin et al., 2017). To test if the pre-formed ERL1-TMM receptor complex is required for the internalization of ERL1, we took advantage of EPFL6, a peptide related to EPF1 with a distinct property (Figure 5) (Abrash and Bergmann, 2010; Abrash et al., 2011). EPFL6 is normally expressed in the internal tissues of hypocotyls and stems, but not in the stomatal-lineage cells. Unlike EPF1, ectopic EPFL6 is a potent inhibitor of stomatal development, even in the *tmm* mutant background (Figure 5A, B) (Abrash and Bergmann, 2010; Abrash et al., 2011; Uchida et al., 2012). Using a similar strategy as MEPF1, we purified biologically active, predicted mature EPFL6 (MEPFL6) peptide. Indeed, the inhibition of stomatal formation by MEPFL6 is more sensitive in *tmm* mutant than in wild type (Figure S4). In contrast to MEPF1, MEPFL6 application induced ERL1-YFP internalization in a dosage-dependent manner regardless of the presence or absence of TMM (Figure 5C-F). The results indicate that TMM is not required for EPFL6-triggered ERL1-YFP internalization. Rather, the ERL1-YFP endocytosis accurately reflects the activity of ERL1 signaling to inhibit stomatal development (Figure 5 and Figure S4), thereby supporting the notion that distinct EPF/EPFL peptide ligands activate a sub-population of ERL1 receptor complexes to internalize through a TMM-based discriminatory mechanism.

**Figure 5.**
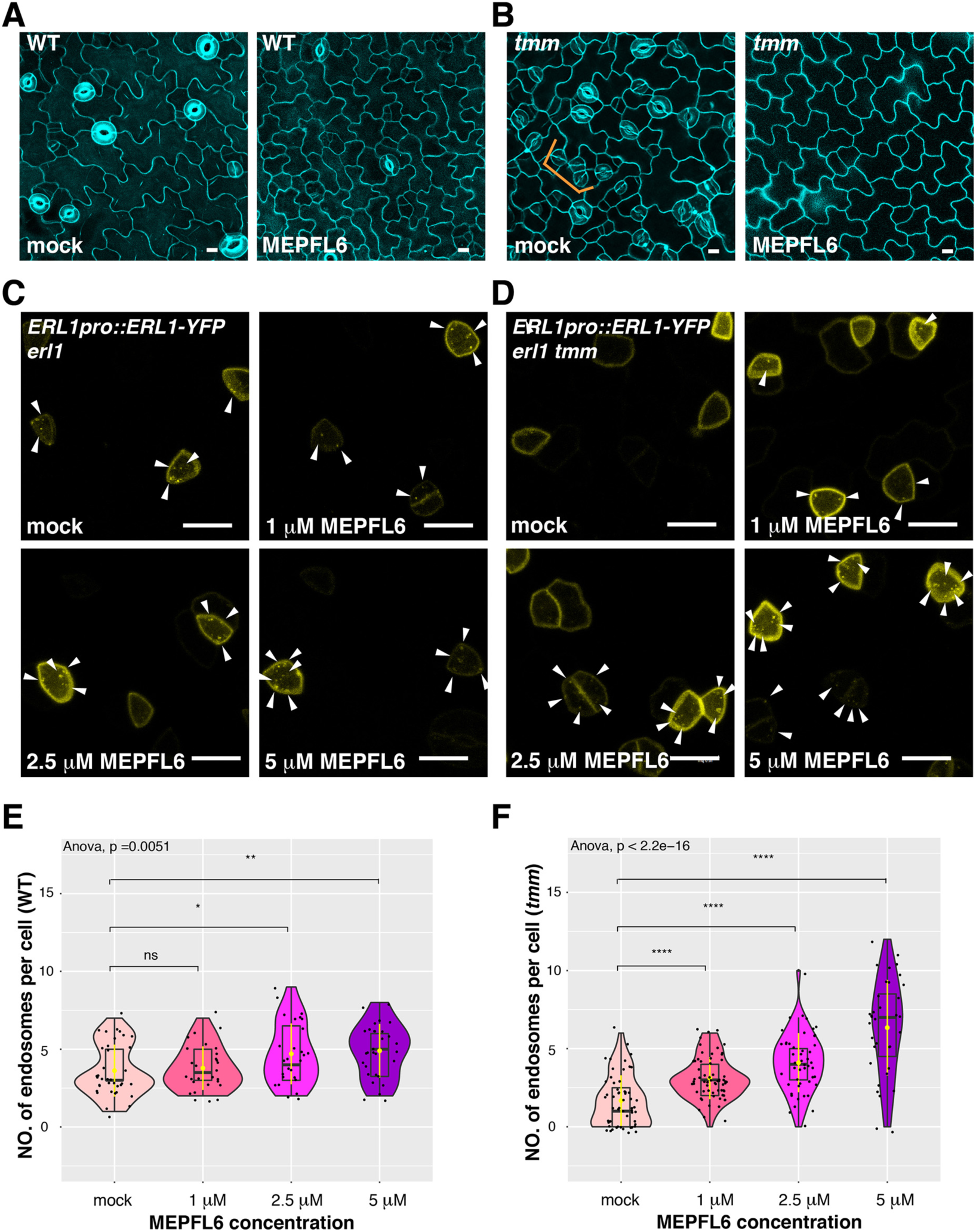
MEPFL6 triggers ERL1-YFP internalization in both *erl1* and *erl1 tmm*. (**A**) Representative confocal microscopy images of cotyledon abaxial epidermis from the 5-day-old wild type seedlings treated with mock (left) or 5 µM MEPFL6 (right). Scale bars =10 µm. (**B**) Shown are representative confocal microscopy images of cotyledon abaxial epidermis from the 5-day-old *tmm* seedlings treated with mock (left) or 5 µM MEPFL6 (right). Scale bar =10 µm. (**C**) Representative images of ERL1-YFP in *erl1* treated with mock (top left), 1 µM MEPFL6 (top right), 2.5 µM MEPFL6 (bottom left) and 5 µM MEPFL6 (bottom right) are shown. Arrowheads indicate endosomes. Scale bar = 10 µm. (**D**) Representative images of ERL1-YFP in *erl1 tmm* treated with mock (top left), 1 µM MEPFL6 (top right), 2.5 µM MEPFL6 (bottom left) and 5 µM MEPFL6 (bottom right) are shown. Arrowheads indicate endosomes. Scale bars = 10 µm. (**E**) Quantitative analysis of the number of ERL1-YFP-positive endosomes per cell at different concentrations of MEPFL6 application in *erl1* shown as a Violin plot. Median values are shown as lines in the boxplot, and mean values are shown as yellow dots in the plot. Dots, individual data points. ANOVA was performed for comparing all samples, and T-test was performed for a pairwise comparisons of samples treated with the mock and different concentration of MEPFL6. p values were indicated for every pairwise comparison. n= 37, 28, 27, 30 for treatment with mock, 1 µM, 2.5 µM, 5 µM MEPFL6. (**F**) Quantitative analysis of the number of ERL1-YFP-positive endosomes per cell at different concentrations of MEPFL6 application in *erl1 tmm* shown as a Violin plot. Dots, individual data points. Median values are shown as lines in the boxplot, and mean values are shown as yellow dots in the plot. ANOVA was performed for comparing all samples, and T-test was performed for a pairwise comparisons of samples treated with the mock and different concentration of MEPFL6. P values were indicated between every two compared samples. n= 55, 63, 48, 35 for treatment with mock, 1 µM, 2.5 µM, 5 µM MEPFL6.

### An antagonistic EPFL peptide, Stomagen, elicits retention of ERL1-YFP in the endoplasmic reticulum

Stomagen promotes stomatal development by competing with other EPFs for binding to the same receptor complex, including ERL1 (Figure 6A) (Kondo et al., 2010; Sugano et al., 2010; Lee et al., 2015; Lin et al., 2017; Qi et al., 2017). Because the activated ERL1 receptor undergoes endocytosis to MVBs, we sought to address the role of Stomagen on subcellular dynamics of ERL1. For this purpose, we first applied bioactive Stomagen peptide on seedlings expressing *ERL1pro::ERL1-YFP* in *erl1.* Unlike in mock-treated samples, YFP signal was detected inside of the cells (Figure S5A). We subsequently treated Stomagen peptides to ERL1-YFP in the *erecta erl1 erl2* triple mutant background to remove any potential redundancy among three ERECTA-family receptors. Strikingly, strong ERL1-YFP signals were detected in a ring-like structure surrounding the nucleus (Figure 6B), which co-localizes with the endoplasmic reticulum marker protein RFP-KDEL (Figure 6B). Thus, Stomagen application results in accumulation of ERL1 in the endoplasmic reticulum.

**Figure 6.**
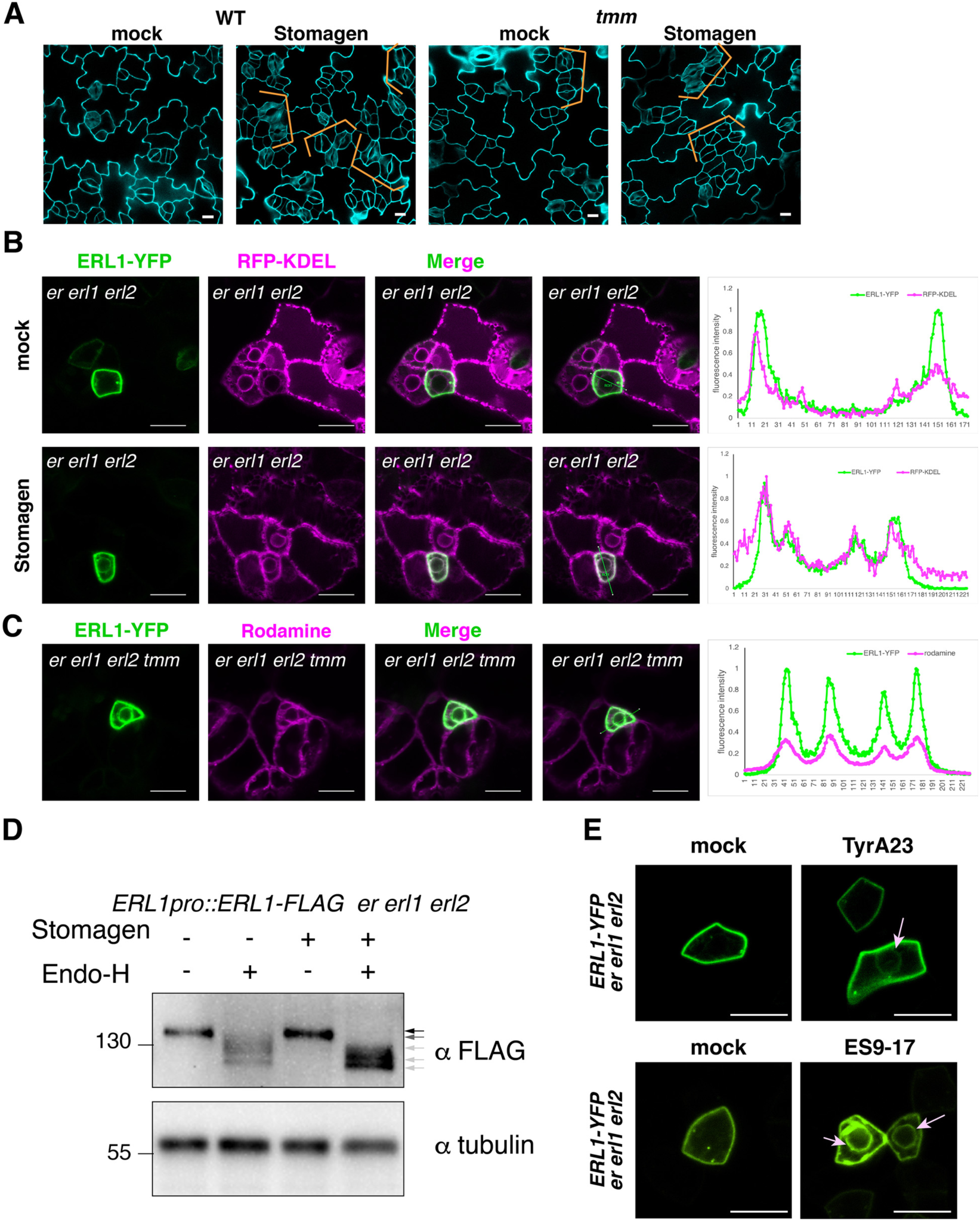
Stomagen application confers accumulation of ERL1 in endoplasmic reticulum. (**A**) Representative confocal microscopy images of cotyledon abaxial epidermis from the 5-day-old wild type seedlings (left two) or *tmm* seedlings (right two) treated with mock (first and third from the left) or 5 µM Stomagen (second and forth from the left). Scale bars = 10 µm. (**B**) Representative confocal microscopy images of ERL1-YFP (left column) and co-localization analysis with the endoplasmic reticulum marker RFP-KDEL (second left column) in the abaxial epidermis of cotyledons of the 5-day-old *erecta* (*er*) *erl1 erl2* seedlings treated with mock (top row) or 5 µM Stomagen (bottom row). Merged images are shown in the third left column. Fourth column shows the line slicing along which quantification analysis of the YFP intensity (green) and RFP intensity (magenta) was done; graphs are shown on the right, with two middle peaks (pointed by arrowheads) showing signals from the endoplasmic reticulum and two big peaks on both sides showing signals of the plasma membrane. Scale bars = 10 µm. (**C**) Representative confocal microscopy images of ERL1-YFP (left) in the abaxial epidermis of cotyledons of the 5-day-old *erecta erl1 erl2* seedlings stained with the endoplasmic reticulum dye Rodamine (second left column). The merged image is shown in the third left column. Quantification analysis of the YFP intensity (green) and RFP intensity (magenta) along the line drawn in the right image is shown as a graph on the right, with two middle peaks (pointed by arrowheads) showing signals from the endoplasmic reticulum and two big peaks on both sides showing signals of the plasma membrane. Scale bars = 10 µm. (**C**) Representative confocal microscopy images of ERL1-YFP (left) in the abaxial epidermis of cotyledons of the 5-day-old *erecta erl1 erl2* seedlings stained with the endoplasmic reticulum dye Rodamine (second left column). The merged image is shown in the third left column. Quantification analysis of the YFP intensity (green) and RFP intensity (magenta) along the line drawn in the right image is shown as a graph on the right, with two middle peaks (pointed by arrowheads) showing signals from the endoplasmic reticulum and two big peaks on both sides showing signals of the plasma membrane. Scale bars = 10 µm. (**D**) Immunoblot analysis of 3-day-old ERL1-FLAG *erecta erl1 erl2* seedlings treated with mock or 5 µM Stomagen for 2 days and then digested without or with Endo-H. Top panel shows the ERL1-FLAG detected by α-FLAG. Lower panel shows the loading control of Tubulin detected by α-Tubulin. Arrows indicate the ERL1 bands detected without or with Endo-H digestion. (**E**) Representative confocal microscopy images of ERL1-YFP expressed in *erecta erl1 erl2* seedlings treated with mock (top left) or 50 µM Tyr A23 (top right); mock (bottom left) or 100 µM ES9-17 (bottom right). Arrow indicates the ring-like structure, characteristics of endoplasmic reticulum localization, detected after treatment with Tyr A23 or ES9-17. Scale bars = 10 µm.

Next, to examine a consequence of inactive ERL1 receptor on its subcellular dynamics, we applied Stomagen peptide on *tmm* seedlings expressing *ERL1pro::ERL1-YFP* and carefully reexamined the inner cellular signal. Very faint ring-like structures were highlighted by ERL1-YFP in both mock and Stomagen-treated meristemoids (Figure S5A). This was enhanced in the *erecta erl1 erl2 tmm* quadruple mutant (Figure 6C). These ERL1-YFP signals co-localized with Rhodamine B hexyl esters, a dye that stains the endoplasmic reticulum (Figure 6C). Thus, in the absence of TMM, ERL1 accumulates in the endoplasmic reticulum.

To biochemically characterize the effects of Stomagen application and *tmm* mutation on ERL1 accumulation in the endoplasmic reticulum, we further performed endoglycosidase H (Endo-H) enzymatic sensitivity assays. Endo-H cleaves N-glycans of proteins in the endoplasm reticulum, including LRR-RLKs (Jin et al., 2007; Nekrasov et al., 2009), but not the remodeled glycan chains of proteins transported to the Golgi or further. To detect slight molecular mass changes, proteins from *erecta erl1 erl2* triple mutant seedlings rescued by *ERL1pro::ERL1-FLAG* were subjected to Endo-H treatment (see Methods). Under normal conditions, ERL1-FLAG is detected as a single band on immunoblots (Figure 6D, black arrow). The Endo-H digestion resulted in a faster mobility of ERL1-FLAG protein with at least three different sizes, suggestive of heterogeneous glycans (Figure 6D, dark gray and light gray arrows). In contrast, ERL1-FLAG protein from Stomagen-treated seedlings was hypersensitive to Endo-H and cleaved completely (Figure 6D, light arrow). Likewise, the *tmm* mutation enhanced the Endo-H sensitivity of ERL1 (Figure S5B), indicating increase in endoplasmic reticulum retention.

Because exogenous application of Stomagen blocks the activation of ERECTA-family signaling (Lee et al., 2015) and results in stomatal clustering (Figure 6A), we sought to address if insufficient internalization of ERL1 from the plasma membrane triggers its stalling in endoplasmic reticulum. For this purpose, we first treated *erecta erl1 erl2* seedlings expressing ERL1-YFP with Tyrphostin A23 (Tyr A23), an inhibitor that has been widely used to block clathrin-mediated endocytosis in plant cells (Banbury et al., 2003; Santuari et al., 2011). Indeed, the TyrA23 treatment enhanced ERL1-YFP signals in the endoplasmic reticulum (Figure 6E, pink arrow), whereas in mock ERL1-YFP was only detected on the plasma membrane and endosomes.

A recent report showed that Tyr A23 functions as a protonophore, which inadvertently blocks endocytosis through cytoplasmic acidification (Dejonghe et al., 2016). Chemical screening and subsequent derivatization identified Endosidin 9-17 (ES9-17) as a specific inhibitor of clathrin-mediated endocytosis without the side effects of cytoplasmic acidification (Dejonghe et al., 2019). We sought to test the effects of ES9-17 on ERL1-YFP subcellular localization to rule out the possibility that retention of ERL1-YFP in the endoplasmic reticulum is due to cellular acidification. ES9-17 previously has been applied only to root cells (Dejonghe et al., 2019). We first optimized the treatment condition for developing seedling shoots (see Methods). At 100 μM, ES9-17 inhibited the internalization of FM4-64 dye in epidermal pavement cells and stomatal-lineage cells, just like in root cells (Figure S6). Under this condition, ES9-17 treatment caused the accumulation of ERL1-YFP in the endoplasmic reticulum, just like the Tyr A23 treatment (Figure 6E). Taken together, our cell biological, pharmacological, and biochemical analyses reveal that inefficient endocytosis due to perception of an antagonistic peptide, Stomagen, as well as loss of co-receptor TMM, causes ERL1-YFP retention in the endoplasmic reticulum.

## Discussion

In this study, we revealed that ERL1 endocytosis accurately reflects EPF signal perception based on three pieces of evidence (Figure 7): first, both EPF1 and EPFL6 peptides trigger ERL1 endocytosis. Second, in the absence of the co-receptor TMM, ERL1 endocytosis is compromised and becomes insensitive to EPF1 application. Third, the kinase domain of ERL1 is required for ERL1 endocytosis. EPF1 and EPFL6 peptide application increased ERL1 population in endosomes in a dosage-dependent manner (Figures 4, 5). ERL1 population in the Wortmannin bodies is reduced in absence of TMM whereas the number of ERL1-marked BFA bodies is not affected (Figures 2, 3), indicating that ERL1 is constitutively recycled whereas the receptor activation triggers endocytosis to MVB, and eventually to a vacuole. In this aspect, the subcellular dynamics of ERL1 resembles that of FLS2, which is also constitutively recycled but rapidly removed from the cell surface upon flg22 perception (Robatzek et al., 2006; Smith et al., 2014). Unlike FLS2, however, a vast majority of ERL1-YFP signal still remained at the plasma membrane even after treatment of 5 µM MEPF1 (Figure 4). These differences could be attributed to the roles of FLS2 and ERL1 in immunity *vs*. development, respectively. FLS2 mediates acute pathogen-induced defense response, whereas ERL1 likely detects endogenous peptides to influence slower processes of cell division and differentiation. A recent study showed, however, that defects in the clathrin-mediated FLS2 endocytosis impair only a subset of FLS2-mediated immune responses (Mbengue et al., 2016). Thus, the precise contributions of endocytosis and cellular response remain open questions. Posttranslational modifications, such as phosphorylation and ubiquitination, of the receptors have emerged as key regulators of receptor subcellular dynamics in FLS2 and BRI1 (Robatzek et al., 2006; Lu et al., 2011; Martins et al., 2015; Zhou et al., 2018). While specific posttranslational modifications of ERL1 are yet unknown, our finding, that dominant-negative ERL1 lacking the entire cytoplasmic domain fails to internalize (Figure 3), suggests that ERL1 phosphorylation may facilitate its endocytosis.

**Figure 7.**
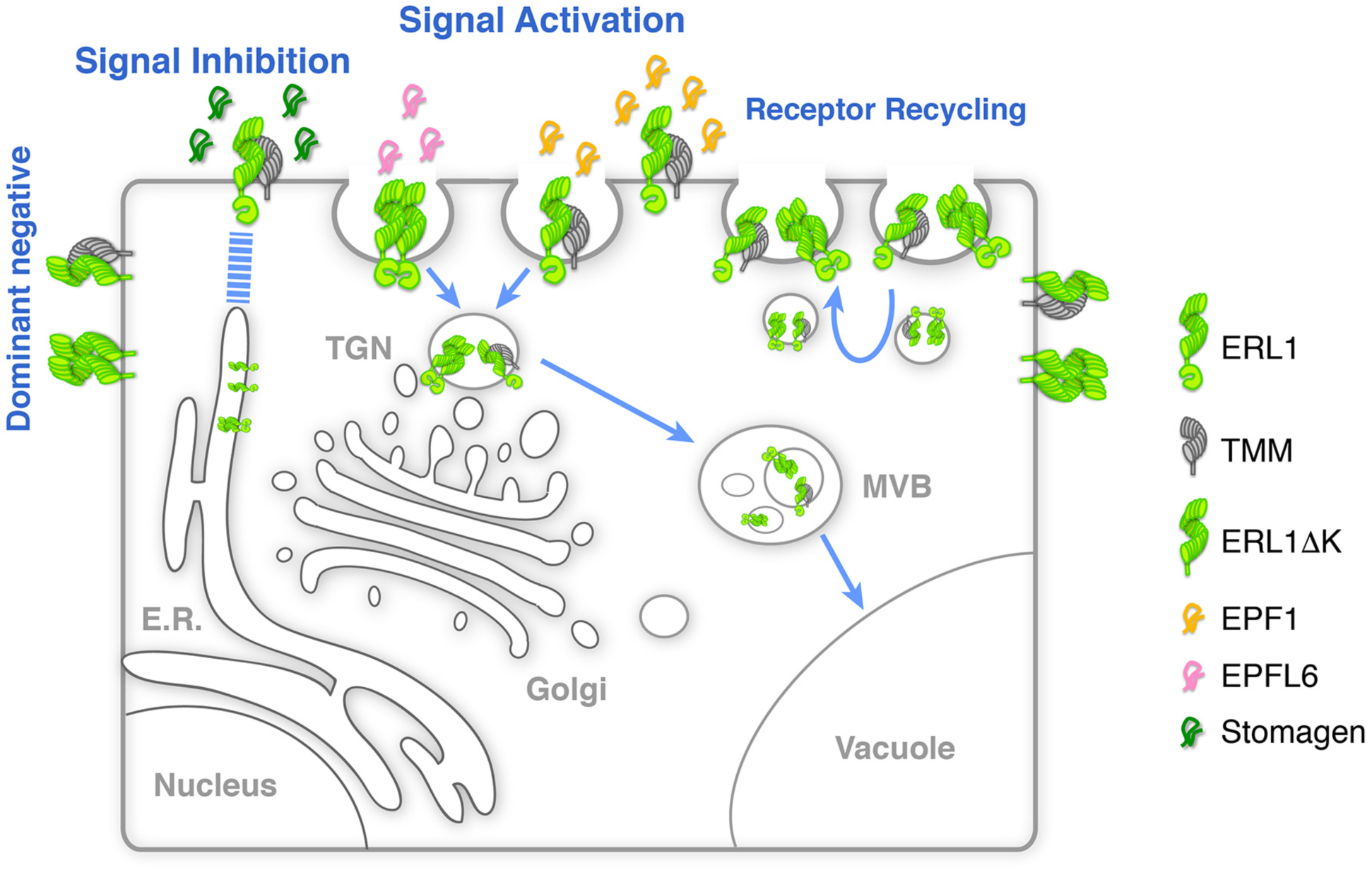
Schematic model of ERL1 subcellular dynamics triggered by diverse EPF peptides with different biological activities. ERL1 (light green) is constitutively recycling and follows BFA-sensitive endosomal pathway (Receptor Recycling). EPF1 (orange) and EPFL6 (pink) peptide ligands both activate ERL1 to inhibit stomatal differentiation, trigger ERL1 trafficking via Wm-sensitive MVB, to vacuole (Signal Activation). EPF1-triggered ERL1 trafficking requires the presence of TMM (gray). In contrast, EPFL6 triggers ERL1 trafficking in TMM-independent manner. Stomagen (dark green), which blocks ERL1 signaling, causes stalling of ERL1 in endoplasmic reticulum (E.R.) (Signal Inhibition). The dominant-negative ERL1ΔK is overwhelmingly plasma-membrane localized, with undetectable level of MVB-mediated internalization (Dominant Negative).

It has been shown that EPF1, but not EPFL6, requires TMM for the inhibition of stomatal development (Hara et al., 2007; Abrash and Bergmann, 2010). Likewise, structural analyses of the EPF-ERECTA family complexes showed that EPF1, but not EPFL6, requires TMM for binding to the ectodomain of ERECTA family receptors (Lin et al., 2017). Here, we demonstrate that TMM is required for endocytosis triggered by EPF1, but not by EPFL6 (Figs. 2, 4 and 5). Thus, at least two populations of ERL1 receptor complexes must be present on the plasma membrane, with and without TMM. Indeed, our FRAP analysis detected the different mobility of these two ERL1 compositions on the plasma membrane (Figure 2). Multiple compositions of receptor complexes have also been reported in CLV3 signaling, where CLV1 homomers, CLV2/CORYNE (CRN) heterodimers and CLV1/CLV2/CRN multimers co-exist on the plasma membrane (Somssich et al., 2015). However, only the microdomain-localized CLV1/CLV2/CRN multimers can perceive the sole ligand CLV3. In the case of BRI1 and FLS2, pre-formed BRI1-BAK1 complex was detected regardless of BRs whereas FLS2 forms FLS2-BAK1 complex upon flg22 application (Bücherl et al., 2013; Somssich et al., 2015). These receptor complexes are spatially separated, even though BRI1 and FLS2 share the same co-receptor BAK1 (Bücherl et al., 2013; Somssich et al., 2015; Bücherl et al., 2017; Hutten et al., 2017). On the contrary, both compositions of ERL1 complexes are ‘functional’ and ligand-inducible, as they can perceive EPF1 or EPFL6, respectively (Figs, 4 and 5). It is possible that the distinct ERL1 receptor complexes reside in different microdomains on the plasma membrane and undergo different trafficking routes upon the correlated ligand perception. EPF1 triggers ERL1 association with BAK1 (Meng et al., 2015). Examining spatiotemporal subcellular dynamics of ERL1 together with TMM and BAK1 at a super resolution scale may reveal the contribution of each receptor complex for specific signal perception and transduction.

Surprisingly, ERL1 is retained in the endoplasmic reticulum when treated with exogenous Stomagen. Extensive studies support that the steady state of a protein in its subcellular compartment is interdependent on the anterograde and retrograde trafficking routes (Brandizzi and Barlowe, 2013). For example, a secretory protein is often retained in the endoplasmic reticulum when the downstream secretion pathway is compromised (Zheng et al., 2005). Blocking the endoplasmic reticulum-to-Golgi retrograde trafficking will accelerate protein transport to the cell surface (Fossati et al., 2014). It is thus possible that Stomagen binding prevents the ERL1 endocytosis and the plasma membrane-accumulated ERL1 interferes with the normal transport of incoming ERL1 from the endoplasmic reticulum. Two additional pieces of evidence support this hypothesis. First, when endocytosis is blocked by Tyr A23 (Banbury et al., 2003) or ES7-19, the improved, specific inhibitor of clathrin heavy chain (Dejonghe et al., 2019), strong ERL1 signals become evident in the endoplasmic reticulum (Figure 6E). Second, in leaves of the *tmm* mutant, where EPFL6 is absent and EPF1-triggered ERL1 endocytosis is compromised, ERL1 also accumulates in the endoplasmic reticulum (Figure 6C and Figure S5). Alternatively, Stomagen-triggered ERL1 accumulation in endoplasmic reticulum may be highlighting the role of the endoplasmic reticulum-plasma membrane contact sites as a direct communication link between the two compartments (Carrasco and Meyer, 2011). The VAP-RELATED SUPPRESSOR OF TMM (VST) family plasma membrane proteins that interact with integral endoplasmic reticulum proteins, have been reported to facilitate ERECTA family-mediated signaling in stomatal development (Ho et al., 2016). Hence, Stomagen perception by the ERL1-TMM complex on the plasma membrane may directly influence signaling via the contact sites and therefore affect the secretion of ERL1 to the cell surface.

Our work revealed the mechanism by which multiple peptide ligands with distinct activities, EPF1, EPFL6, and Stomagen, fine-tune stomatal patterning at the level of the subcellular dynamics of a single receptor, ERL1. Successful development of visible functional peptide ligands and identification of the immediate biochemical events by the ERECTA-family perceiving different EPF peptides will help elucidate the exact roles of receptor trafficking and signaling specifying developmental patterning in plants.

## Acknowledgements

We thank Prof. Takashi Ueda for RFP-Ara7, Syp43-RFP, and Syp22-RFP lines; Prof. Gian Pietro Di Sansebastiano for ST-RFP and KDEL-RFP constructs; Prof. Hugo Zheng for N-ST-YFP and N-YFP-HDEL constructs; Alex Hofstetter for technical assistance of MEPF1 and MEPFL6 peptide purification; Prof. Jenny Russinova for providing ES9-17 and insightful suggestions on experimental designs; Drs. Naoyuki Uchida, Ayami Nakagawa, and Soon-Ki Han for critical comments on the manuscript. This work was supported by the Gordon and Betty Moore Foundation (GBMF3035) to K.U.T., who is a Howard Hughes Medical Institute Investigator.

## Author contributions

Conceived, K.U.T.; Designed experiments, X.Q., K.U.T.; Performed experiments, X.Q.; Peptide refolding and bioassays, X.Q, M.M.; Debugged and ran FrapBot in a local environment, S.Z., K.U.T.; Analyzed data, X.Q., S.Z., K.U.T.; Visualization, X.Q., K.U.T.; Writing-Original Draft, X.Q., K.U.T; Writing-Review & Editing, X.Q., M.M., S.Z., K.U.T.; Project Administration, K.U.T.; Funding Acquisition, K.U.T.

## Methods

### Plant materials and growth conditions

The *Arabidopsis* accession Columbia (Col) was used as wild type. The following mutants and reporter transgenic plant lines used in this study were reported previously: *erecta (er-105)* (Shpak et al., 2005)*; erl1-2 (Shpak et al., 2005); erl2-1* (Shpak et al., 2005)*; epf1-1* (Hara et al., 2007); *tmm-KO* (Hara et al., 2007); *ERL1pro::ERL1-YFP* in *erl1-2, ERL1pro:: ERL1-FLAG in erl1-2* and *erecta erl1-2 erl2-1,* and *ERL1pro::ERL1ΔKinase* in *erl1-2* (Lee et al., 2012); *MUTEpro::ERL1-YFP* in *er-105 erl1-2 erl2-1* and iEPF1 lines (Qi et al., 2017). Transgenic *Arabidopsis* lines expressing *ARA7pro::mRFP-ARA7*, *SYP22pro::mRFP-SYP22*, and *SYP43pro::mRFP-SYP43* are a gift from Prof. Takashi Ueda (NIBB, Japan). *ST-RFP* and *KDEL-RFP* constructs are from Prof. Gian Pietro Di Sansebastiano (Univ. of Salento, Italy). Reporter lines were introduced into respective mutant backgrounds by genetic crosses or by *Agrobacterium*-mediated floral-dipping transformation, and genotypes were confirmed by PCR. Seedlings and plants were grown as described previously (Lee et al., 2012). For a list of PCR-based genotyping primer sequence, see Table S1.

### Recombinant peptide production

Expression, purification, and refolding of predicted mature EPF1 (MEPF1) or EPFL6 (MEPFL6) peptides were performed as described previously (Lee et al., 2012), except for the following. His-tagged MEPF1 or MEPFL6 was affinity purified on 5 ml His-Trap HP column (GE Healthcare) using NGC^™^ Chromatography System (Bio-Rad). Inclusion bodies from 1.0 L of *E. coli* were solubilized in guanidine hydrochloride (Gdn-HCl) buffer (6.0 M Gdn-HCl, 500 mM NaCl, 5 mM imidazole, 1 mM 2-mercaptoethanol, 50 mM Tris, pH 8.0) and loaded onto the column and washed with 10 column volumes (50 mL) of Wash Buffer (8.0 M urea, 500 mM NaCl, 30 mM imidazole, 1 mM β-mercaptoethanol, 50 mM Tris, pH 8.0) at a flow rate of 3.00 ml/min, and MEPF1 or MEPFL6 peptides were eluted with a 0-100 % gradient of Wash to Elution Buffer (8.0 M urea, 500 mM NaCl, 500 mM imidazole, 1 mM β-mercaptoethanol, 50 mM Tris, pH 8.0) over 10 column volumes at 3.00 mL/min prior to refolding. The quality of refolded peptide was analyzed by HPLC (Walters DataPrep 300), its bioactivity was confirmed using Arabidopsis seedlings, and bioassay on Arabidopsis seedlings were performed as described previously (Lee et al., 2012; Lee et al., 2015). For the dose-response analysis of EPFL6, the R-package ‘drc’ (Ritz et al., 2015) was used to fit the binding curve to the generalized log logistic distribution (Uchida et al., 2018).

### Pharmacological treatment

BFA (Sigma: Cat No. B7651) and Wortmannin (Sigma: Cat No. W1628) were dissolved as 10 mM stock using ethanol and DMSO, respectively. For BFA treatment, cotyledons of 7-day-old seedlings were removed, and the rest of the seedlings were immersed into either mock (0.3% of ethanol), or 30µM BFA solution, vacuumed for 1min, and immersed for 30 min before imaging. For Wortmannin treatment, seedlings were treated with 25 µM Wortmannin in 0.25% DMSO. 0.25% DMSO solution was used as a mock condition. For MEPF1 and MEPFL6 treatment, purified peptide solution was diluted to 5 µM using liquid ½ MS media. Cotyledons of 7-day-old seedlings were removed, and the rest of the seedlings were immersed into the above solutions, vacuumed for 1 min, and immersed for 10 min before imaging. The same procedure was done for Stomagen treatment except that the seedlings were immersed into the solution for 1 hour. For co-treatment of cycloheximide (CHX: Sigma, C4859) and BFA, 7-day-old seedlings, with cotyledons moved, were immersed into 50 µM CHX for 1 hour followed by either mock (0.3% of ethanol), or 30µM BFA solution, vacuumed for 1min, and immersed for 30 min before imaging.

For Tyrphostin A23 (Sigma: Cat No. T7165) treatment, Tyrphostin A23 was dissolved as 50 mM stock using DMSO. 5-day-old seedlings were immersed into either mock (0.1% of DMSO) or 50 µM Tyrphostin A23 solution, vacuumed for 1min, and immersed for 1 hour before imaging. ES9-17 was generously provided by Dr. Eugenia Russinova (VIB, Gent). As suggested, ES9-17 was dissolved as 50 mM stock using DMSO. For ES9-17 and FM 4-64 treatment on true leaves, cotyledons of 7-day-old seedlings were removed, and the rest of the seedlings were immersed into either mock (1/2 MS medium with 0.4% of DMSO) or ES9-17 solution (1/2 MS medium with 50 µM ES9-17) followed by 5 µM FM 4-64 (Thermo Fisher, T13320) staining for 30 min before imaging. For ES9-17 and FM 4-64 treatment in roots, 3-day-old seedlings were immersed into either mock (1/2 MS medium with 0.4% of DMSO), or ES9-17 solution (1/2 MS medium with 100 µM ES9-17), followed by FM 4-64 (5 µM) staining for 30 min before imaging.

For Rhodamine B (Sigma: Cat No. R6626) hexyl ester treatment, Rhodamine B hexyl ester was dissolved as 16mM stock using DMSO. 5-day-old seedlings were immersed into either mock (1% of DMSO) or 160 µM Rhodamine B hexyl ester solution for 30 min before imaging.

### Protein extraction, enzymatic assay (Endo-H), and protein gel immunoblot analysis

For Endo-H (NEB: Cat No. P0703S) assays, *erecta erl1 erl2* seedlings with functional *ERL1pro::ERL1-FLAG* were grown on ½ MS media plates for 3 days and then transferred to ½ MS liquid media with either Tris-HCl buffer (pH 8.8) or 5 µM Stomagen peptide in a 24-well cluster plate at room temperature for one day before being pooled for harvest. Plant materials were ground in liquid nitrogen, and then extracted with buffer (100 mM Tris-HCl pH 8.8, 150 mM NaCl, 1 mM EDTA, 20% glycerol, 20 mM NaF, 2 mM Na_3_VO_4_, 1 mM PMSF, 1% Triton X-100, 1 tablet per 50 ml extraction buffer of complete^TM^ proteinase inhibitor cocktail, Roche). The extracts were briefly sonicated at 4 °C and centrifuged at 4,000 r.p.m. for 10min at 4 °C to remove cell debris. The supernatant was then ultracentrifuged at 100,000g for 30min at 4 °C. Total protein concentration was determined using a Bradford assay (Bio-Rad: Cat No. 5000006) before adjustment. The solution was incubated with Dynabeads Protein G (Invitrogen: Cat No. 10004D) conjugated with mouse monoclonal anti-FLAG M2 (Sigma: Cat No. F-3165) for 2 hours with slow rotation at 4 °C, followed by washing with TBS with 0.1% Tween 20. The immunoprecipitates were eluted with 2x SDS sample buffer (100 mM Tris-HCl at pH 6.8, 4% SDS, 0.02% Bromophenol Blue, 20% glycerol, 2% 2-mercaptoethanol, 1% proteinase inhibitor cocktail) by boiling for 10 min. Each immunoprecipitate was then separated into two aliquots, treated with either water or Endo-H for 10min at 37 °C. Immunoblot analysis was performed using mouse monoclonal anti-FLAG M2 (Sigma: Cat No. F-3165; 1:5,000) antibody as primary antibody, and horseradish peroxidase-conjugated goat anti-mouse IgG (GE Healthcare: Cat No. NA931VS; 1:50,000) as secondary antibody. For loading control, immunoblot was performed using mouse anti-tubulin (Millipore Sigma: Cat No. MABT205; 1:5000). The protein blots were visualized using Chemiluminescence assay kit (Thermo Scientific: Cat No. 34095).

### Confocal microscopy and image analysis

Confocal microscopy images were taken on the Leica SP5X-WLL inverted confocal microscope (Solms, Germany). Time-lapse imaging of ERL1-YFP true leaves was prepared as described previously (Peterson and Torii, 2012). ERL1-YFP internalization imaging was done with a 63x/1.2 W Corr lens on Leica SP5X. 514 nm laser was used to excite YFP and emission window of 518-600 nm was used to collect YFP signal. For the multicolor images of YFP and RFP, true leaves of 7-day-old transgenic seedlings were observed with a 63x/1.2 W Corr lens on Leica SP5X. 514nm laser was used to excite YFP and 555nm laser was used to excite RFP and FM4-64. Emission filter was set as 518nm-550nm for YFP and 573-630 for RFP and FM4-64. Each experiment was repeated at least three times, each with multiple seedlings. The Leica LAS AF software (http://www.leica-microsystems.com) and Imaris 8.1 (Bitplane) were used for post-acquisition image processing.

### Fluorescence Recovery After Photobleaching (FRAP) analysis

The FRAP experiments were conducted on ERL1-YFP using a 63x/1.2 W Corr lens on the Leica SP5X confocal microscope by photobleaching ∼10% of the plasma membrane with 100% 405 nm laser power. 514 nm laser was used to excite YFP and emission window of 518-600 was used to collect YFP signal. Recovery of fluorescence was monitored in the photobleached plasma membrane for 6 min with 3-second intervals. A non-photobleached region was monitored meanwhile as an internal control. Average intensities of the region of interest were quantified with Leica LAS AF software. The exported data was analyzed and modeled by using the R-based FrapBot software (www.frapbot.kohze.com) (Kohze et al., 2017) with some modification to run on the local lab computer. The FRAP recovery curves were fitted to a single-parameter exponential model to determine the half time.

### Data plots and Statistics

Graphs were generated using R ggplot2. For box plots and violin plots, individual data points are plotted as dot plots. For the violin plots with large sample numbers, the dot plots were jittered with a position of 0.2. All statistical analyses were performed using R. All codes are available upon request.

---

**Figure S1:**
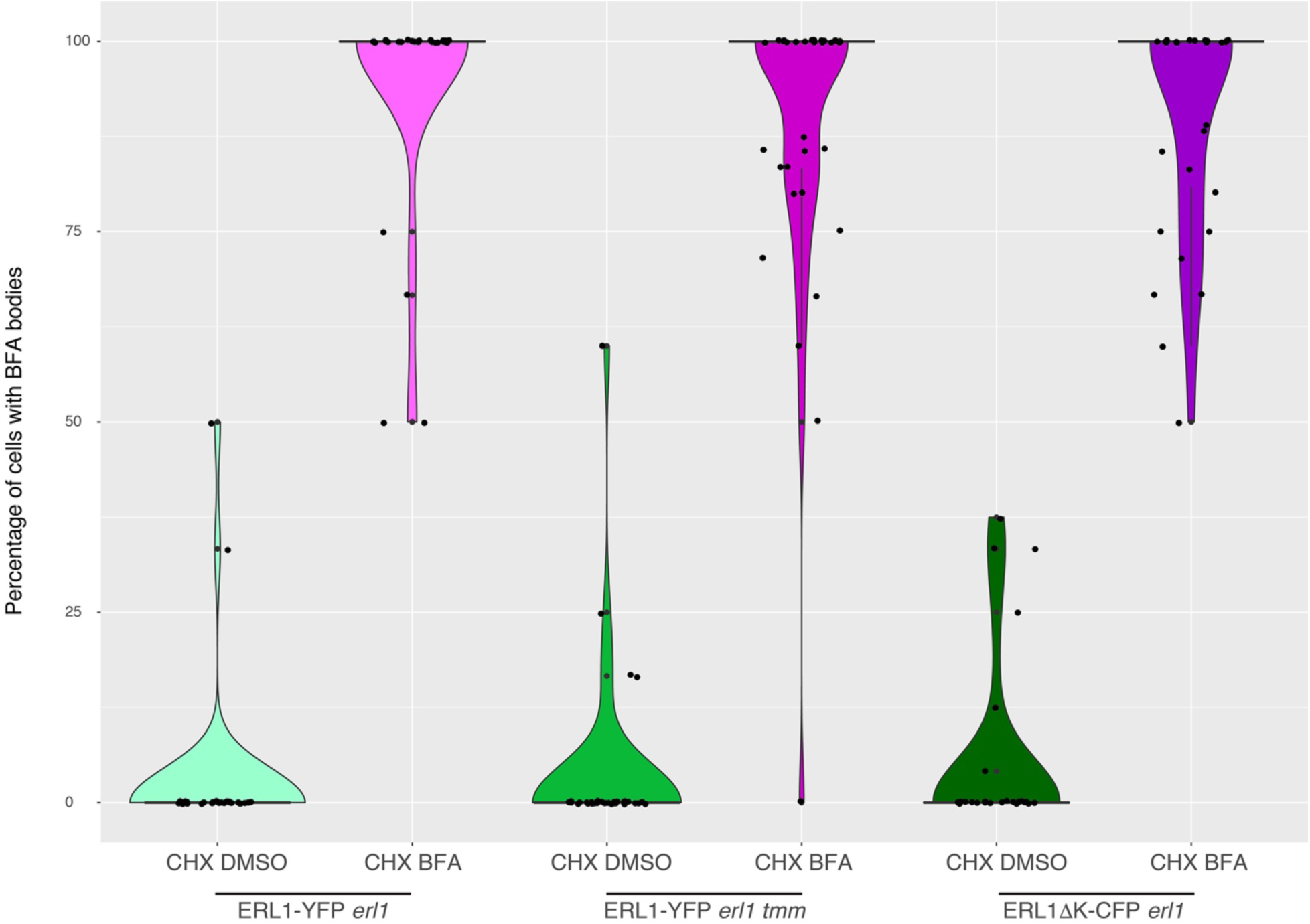
ERL1 BFA body formation in wild-type, *tmm*, or dominant-negative ERL1 background in the presence of cycloheximide. True leaves of 7-day-old Arabidopsis seedlings expressing either ERL1-YFP or ERL1ΔK-CFP were pretreated with 50 µM CHX for 1 hr, followed by treatment with either ethanol or 50 µM BFA for 30 min. Violin plot with boxplot is used to show the quantification of the number of cells with BFA bodies. Individual data points are dot-plotted with jitter. Median values are shown as lines in the boxplot. n= 33 for ERL1-YFP CHX DMSO, n= 29 for ERL1-YFP CHX BFA, n= 34 for ERL1-YFP *tmm* CHX DMSO, n= 34 for ERL1-YFP *tmm* CHX BFA, n= 29 for ERL1ΔK-CFP CHX DMSO, n= 30 for ERL1ΔK-CFP CHX BFA.

**Figure S2:**
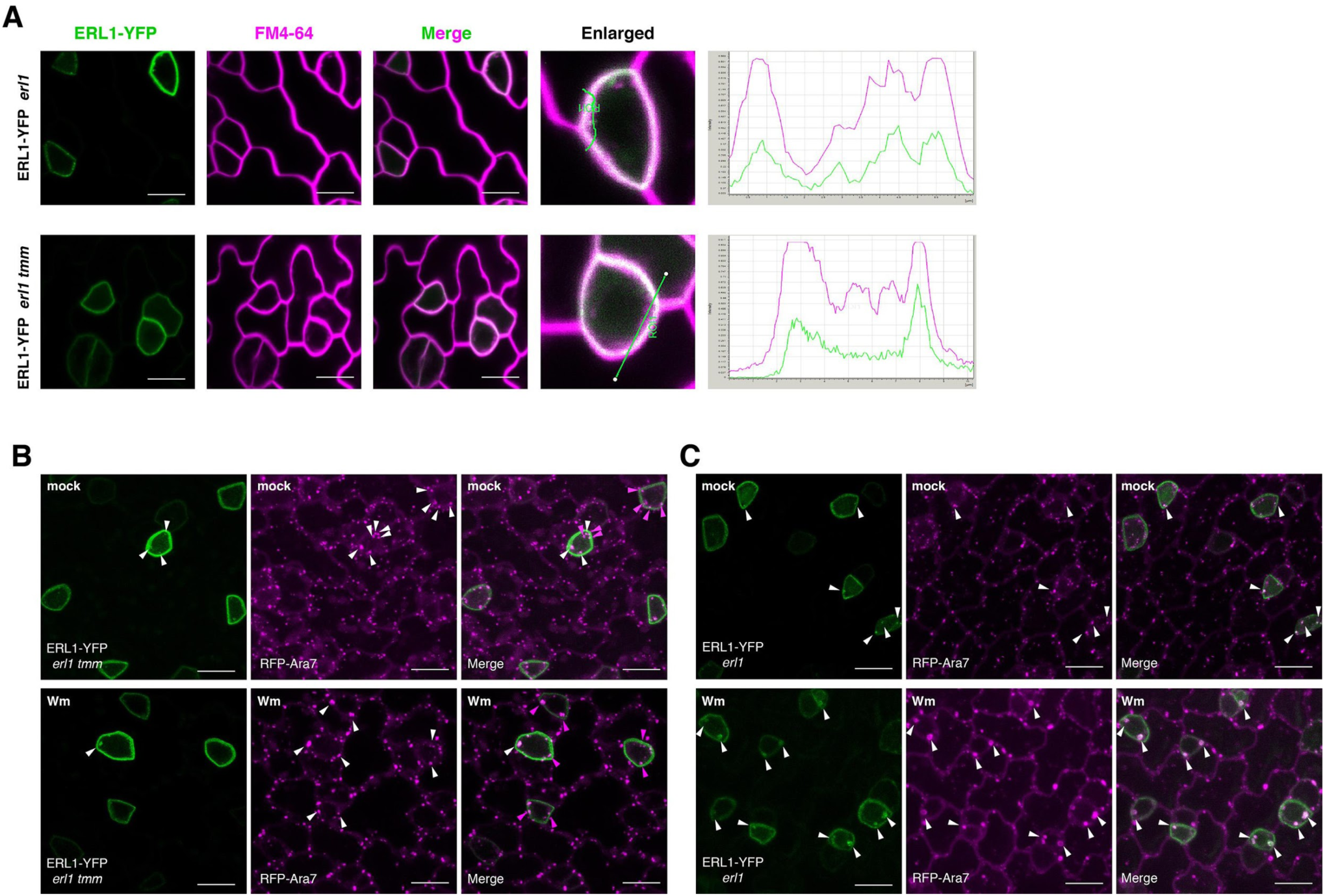
*tmm* mutation does not affect general internalization and the MVB structure. (**A**) Representative confocal microscopy images of ERL1-YFP (green) and FM4-64 staining (magenta) of developing true leaf abaxial epidermis from 7-day-old seedlings expressing *ERL1pro::ERL1-YFP* in *erl1* (top row) and *erl1 tmm* (bottom row). Merged images of ERl1-YFP and FM4-64 staining are shown in the third column from left, and enlarged cells are shown in the last column. Right, fluorescence intensity quantified along the line from different channels showing their co-localization. Scale bars = 10 µm. (**B**) Representative confocal microscopy images of ERL1-YFP (green) and RFP-Ara7 (magenta) of developing true leaf abaxial epidermis from 7-day-old seedlings expressing *ERL1pro::ERL1-YFP* in *erl1 tmm* treated with mock (top row) or with 25 µM Wm (bottom row). Arrowheads indicate endosomes or Wm bodies. Scale bars = 10 µm. (**C**) Shown are representative images of ERL1-YFP (first column) and RFP-Ara7 (second column) in using developing true leaf abaxial epidermis of the 7-day-old seedlings of *erl1* treated with mock/DMSO (top row) or with 25 µM Wm (bottom row). Arrowheads indicate endosomes or Wm bodies. Scale bar = 10 µm.

**Figure S3:**
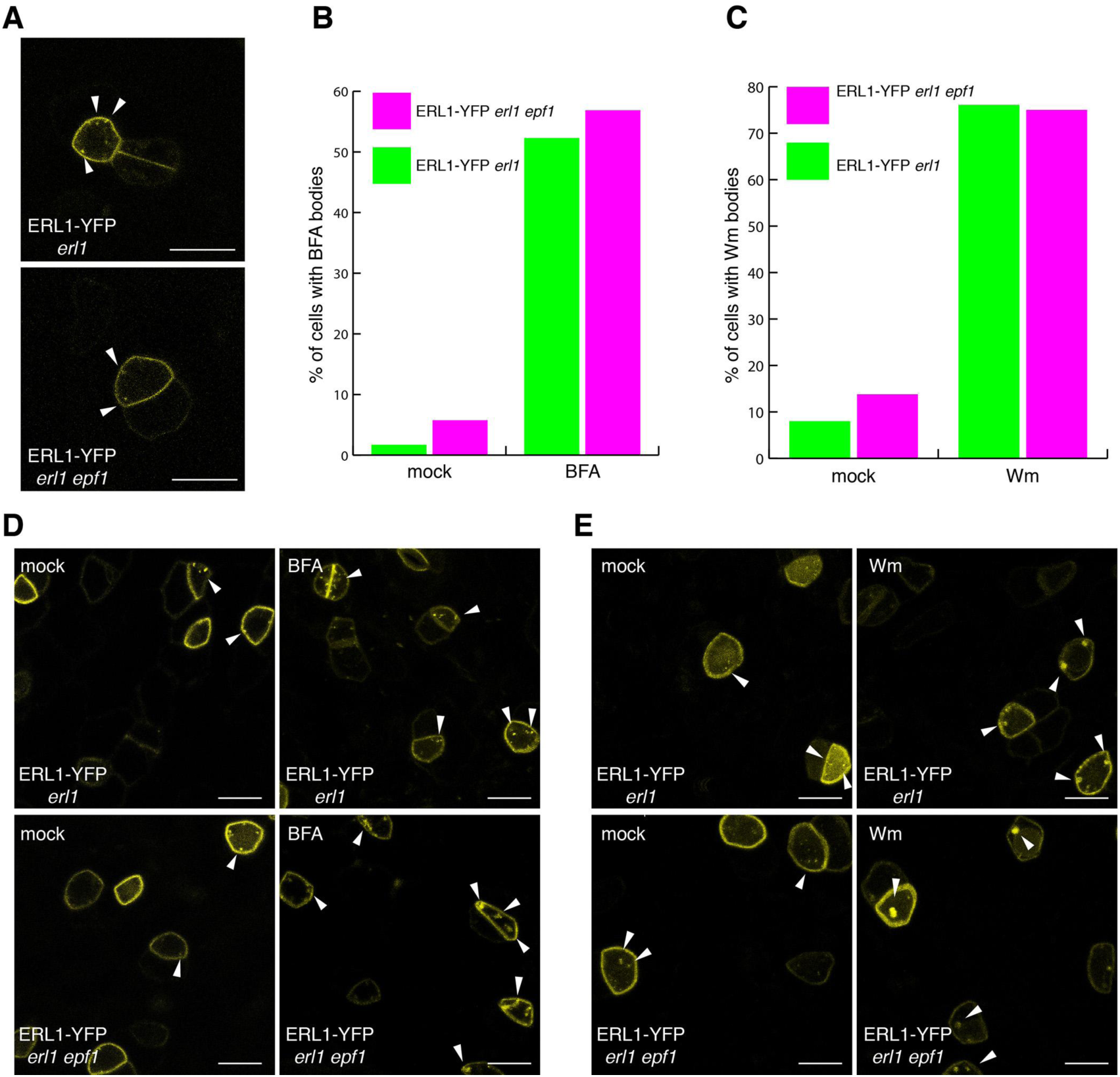
Absence of endogenous EPF1 does not affect ERL1-YFP internalization. (**A**) Representative confocal microscopy images of an abaxial true leaf epidermis from 7-day-old seedling expressing *ERL1pro::ERL1-YFP* in *erl1* (top) and in *erl1 epf1* (bottom); Arrowheads indicate endosomes. Scale bars =10 µm. (**B**) Quantitative analysis of the percentage of cells with BFA bodies when ERL1-YFP in *erl1* (green) or *erl1 epf1* (purple) are treated with mock or 30 µM BFA. n = 117, 139, 67, 51 for mock (*erl1*), mock (*erl1 epf1*), BFA (*erl1*), BFA (*erl1 epf1*). (**C**) Quantitative analysis of the percentage of cells with Wm bodies when ERL1-YFP in *erl1* (green) or *erl1 epf1* (purple) are treated with mock or 25 µM Wm. n = 50, 39, 46, 60 for mock (*erl1*), mock (*erl1 epf1*), Wm (*erl1*), Wm (*erl1 epf1*). (**D**) Representative confocal microscopy images of ERL1-YFP in *erl1* (top row) or in *erl1 epf1* (bottom row) treated with mock (left column) or 30 µM BFA (right column). Arrowheads indicate endosomes or BFA bodies. Scale bars =10 µm. (**E**) Representative confocal microscopy images of ERL1-YFP in *erl1* (top row) or in *erl1 epf1* (bottom row) treated with mock (left column) or 25 µM Wm (right column). Arrowheads indicate endosomes or Wm bodies. Scale bars =10 µm.

**Figure S4:**
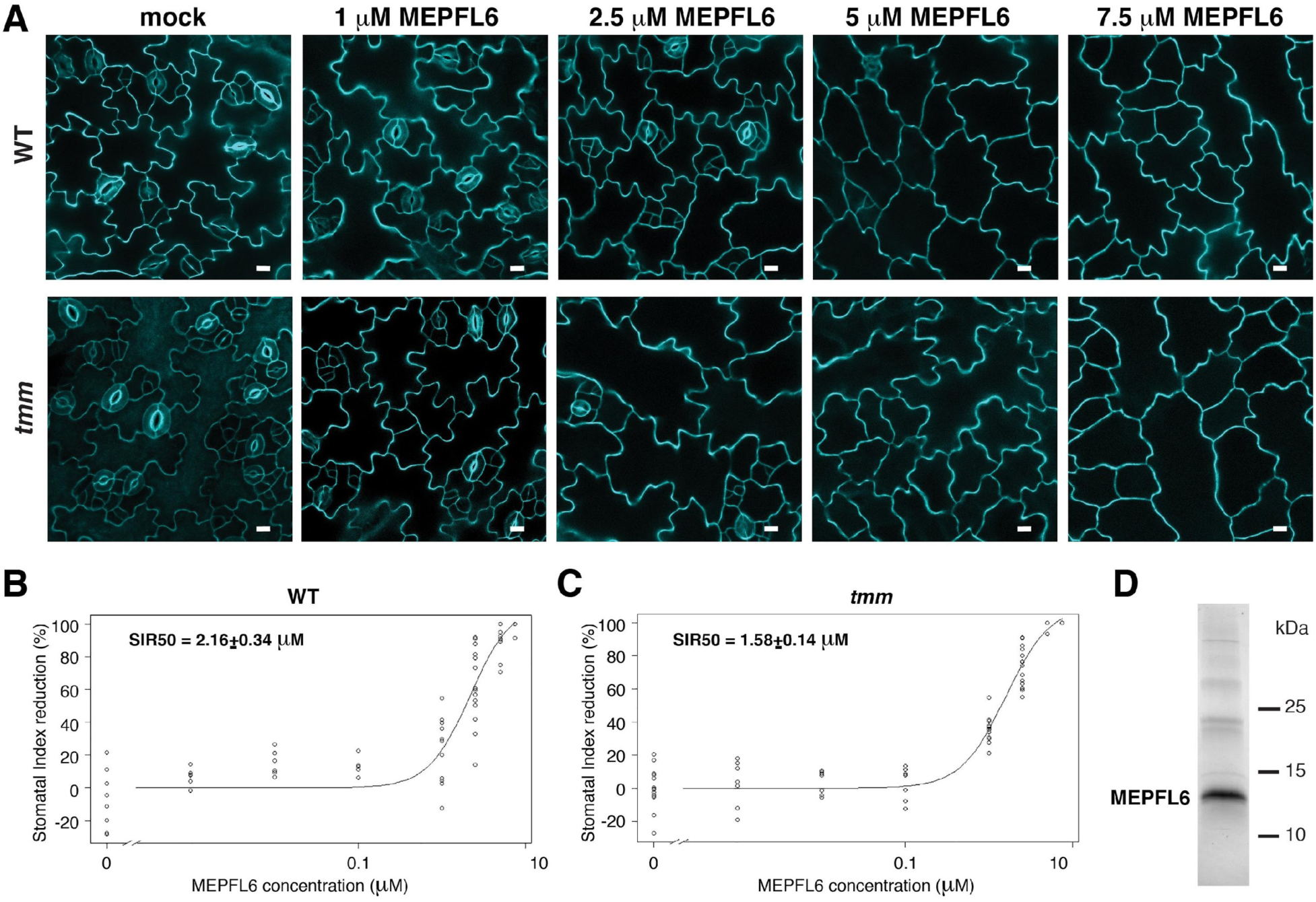
Stomatal development in *tmm* is more sensitive than in wild type to MEPFL6 application. (**A**) Effects of recombinant MEPFL6. Shown are representative confocal images of abaxial epidermis from 5-day-old wild-type (top row) and *tmm* mutant (bottom row) seedlings with mock (left) and increasing concentrations of MEPFL6 (corresponding concentrations are indicated on top of each column). Scale bars = 10 µm. (**B**) Dose response curve of abaxial epidermis stomatal index reduction in 5-day-old wild-type seedlings to different concentrations of MEPFL6. SIR50 indicates the MEPFL6 concentration that causes 50% of **S**tomatal **I**ndex **R**eduction. (**C**) Dose response curve of abaxial epidermis stomatal index reduction in 5-day-old *tmm* mutant seedlings to different concentrations of MEPFL6. SIR50 indicates the MEPFL6 concentration that causes 50% of **S**tomatal **I**ndex **R**eduction. (**D**) SDS-PAGE analysis of predicted MEPFL6-6xHis recombinant protein expressed and purified from *E.coli*.

**Figure S5:**
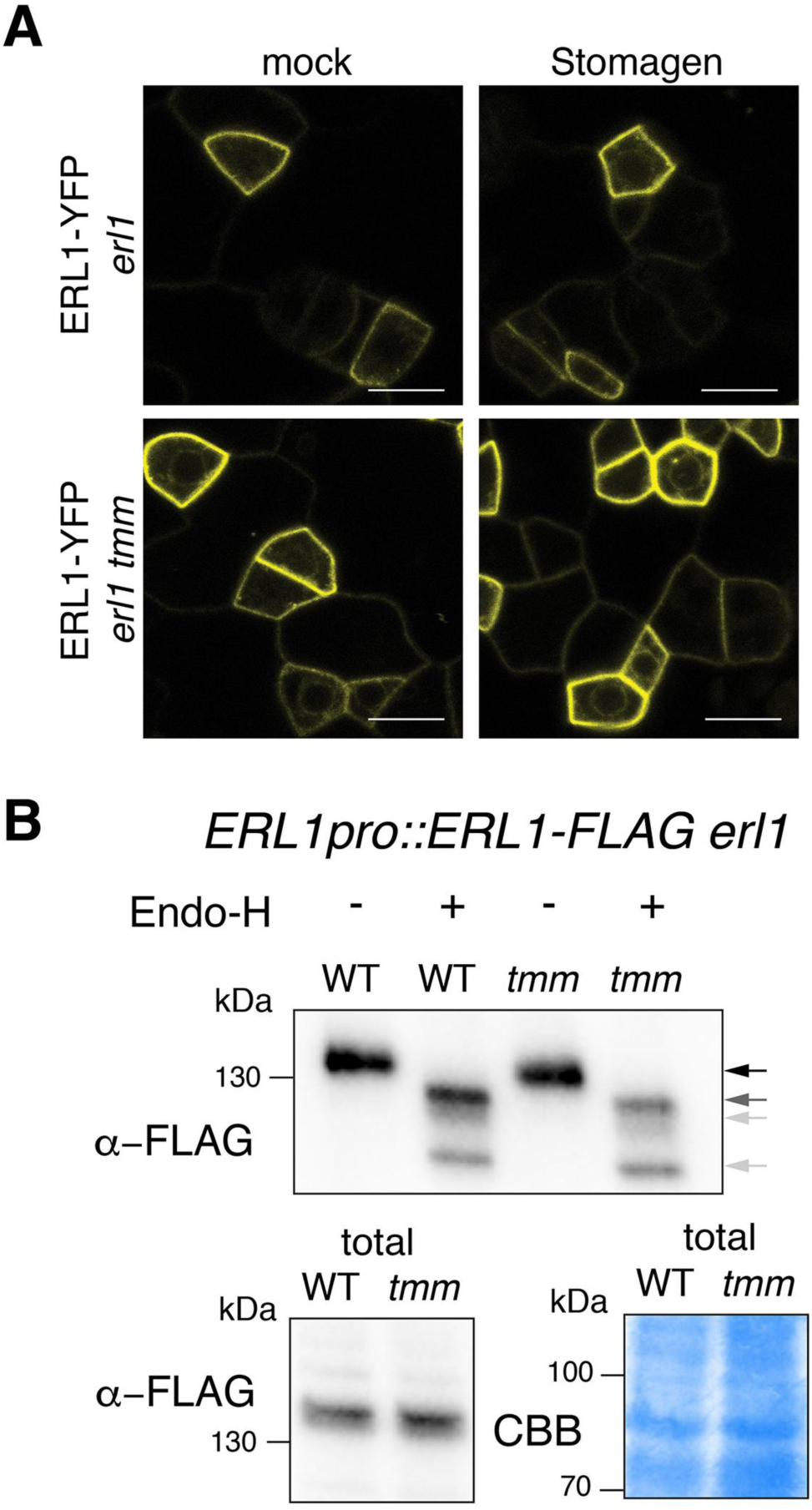
Inefficient endocytosis causes ERL1-YFP to stall in the endoplasmic reticulum. (**A**) Representative confocal microscopy images of ERL1-YFP in the abaxial epidermis of true leaves of the 7-day-old *erl1* seedlings (top row) or *erl1 tmm* seedlings (bottom row) treated with mock (left column) or 5 µM Stomagen (right column). Scale bars = 10 µm. (**B**) Immunoblot analysis of 7-day-old *ERL1pro::ERL1-FLAG erl1* seedlings and *ERL1pro::ERL1-FLAG erl1 tmm* seedlings digested without or with Endo-H. Top panel shows the immunoprecipitated ERL1-FLAG without or with Endo-H digestion detected by α-FLAG. Bottom panels show the ERL1-FLAG from the total protein immune-detected by α-FLAG (left) and those detected by Commassie Brilliant Blue (CBB) staining as a loading control (right). Arrows indicate the ERL1 bands detected without (black arrow) or with (gray arrows) Endo-H digestion.

**Figure S6:**
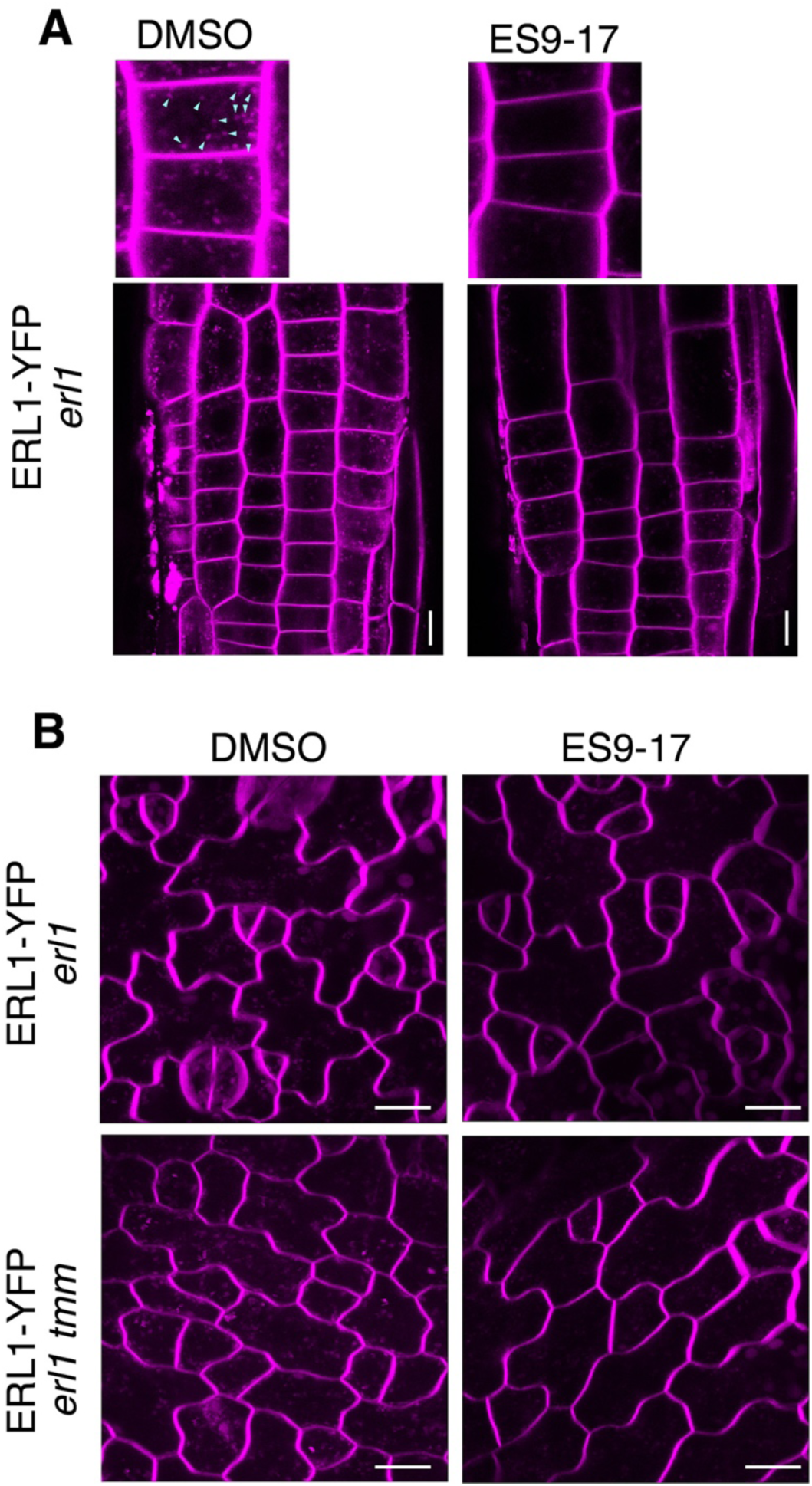
ES9-17 inhibits endocytosis of leaf epidermal cells. (**A**) Primary roots of 3-day-old *erl1-2* seedlings expressing ERL1-YFP treated with DMSO (left) or ES9-17 (right) followed by FM4-64 staining. Top panels, magnified images. Only red channel was used to image FM4-64. The ES7-19 treatment diminishes endocytosis (arrowheads in DMSO). Scale bar = 10 µm. (**B**) True leaves of 7-day-old *erl1* (top) or *erl1 tmm* (bottom) seedlings expressing ERL1-YFP treated with DMSO (left) or ES 9-17 (right) followed by FM4-64 staining. Only red channel was used to image FM4-64. The ES7-19 treatment diminishes endocytosis. Scale bars = 10 µm.

